# Pattern completion and disruption characterize contextual modulation in the visual cortex

**DOI:** 10.1101/2023.03.13.532473

**Authors:** Jiakun Fu, Suhas Shrinivasan, Luca Baroni, Zhuokun Ding, Paul G. Fahey, Paweł A. Pierzchlewicz, Kayla Ponder, Rachel Froebe, Lydia Ntanavara, Taliah Muhammad, Konstantin F. Willeke, Eric Wang, Zhiwei Ding, Dat T. Tran, Stelios Papadopoulos, Saumil Patel, Jacob Reimer, Alexander S. Ecker, Xaq Pitkow, Jan Antolik, Fabian H. Sinz, Ralf M. Häfner, Andreas S. Tolias, Katrin Franke

## Abstract

Vision is fundamentally context-dependent, with neuronal responses influenced not just by local features but also by surrounding contextual information. In the visual cortex, studies using simple grating stimuli indicate that congruent stimuli—where the center and surround share the same orientation—are more inhibitory than when orientations are orthogonal, potentially serving redundancy reduction and predictive coding. Understanding these center-surround interactions in relation to natural image statistics is challenging due to the high dimensionality of the stimulus space, yet crucial for deciphering the neuronal code of real-world sensory processing. Utilizing large-scale recordings from mouse V1, we trained convolutional neural networks (CNNs) to predict and synthesize surround patterns that either optimally suppressed or enhanced responses to center stimuli, confirmed by *in v ivo* experiments. Contrary to the notion that congruent stimuli are suppressive, we found that surrounds that completed patterns based on natural image statistics were facilitatory, while disruptive surrounds were suppressive. Applying our CNN image synthesis method in macaque V1, we discovered that pattern completion within the near surround occurred more frequently with excitatory than with inhibitory surrounds, suggesting that our results in mice are conserved in macaques. Further, experiments and model analyses confirmed previous studies reporting the opposite effect with grating stimuli in both species. Using the MICrONS functional connectomics dataset, we observed that neurons with similar feature selectivity formed excitatory connections regardless of their receptive field overlap, aligning with the pattern completion phenomenon observed for excitatory surrounds. Finally, our empirical results emerged in a normative model of perception implementing Bayesian inference, where neuronal responses are modulated by prior knowledge of natural scene statistics. In summary, our findings identify a novel relationship between contextual information and natural scene statistics and provide evidence for a role of contextual modulation in hierarchical inference.

## Introduction

Across animal species, the processing of sensory information is context-dependent, which can result in varied perceptions of the identical stimulus under different conditions. This adaptive mechanism allows for the flexible adjustment of sensory processing to changing environments and tasks. In the domain of vision, the context is often defined by the global attributes of the broader visual scene. For instance, effective object detection relies not only on the integration of local object features such as contours or textures but also on the visual context surrounding the object (Biederman et al., 1982; Hock et al., 1974). Physiologically, the responses of visual neurons to stimuli within their center receptive field (RF) –— termed the classical RF –— are influenced by stimuli in their surround RF, known as the extra-classical RF. This center-surround contextual modulation is evident across various levels of the visual system, ranging from the retina to the visual cortex (Mcilwain, 1964; Solomon et al., 2002; Hubel and Wiesel, 1965; Knierim and Van Essen, 1992; Keller et al., 2020b; Jones et al., 2012; Rossi et al., 2001; Vinje and Gallant, 2000; Angelucci et al., 2017; Polat et al., 1998; Nurminen and Angelucci, 2014). It is thought to be mediated by both lateral interactions and feedback from higher visual areas (Nassi et al., 2013; Nurminen et al., 2018; Keller et al., 2020a; Adesnik et al., 2012; Angelucci et al., 2017).

Research on how context modulates visual activity has predominantly been using experimental settings with well-defined parametric stimuli, such as oriented gratings. These studies, primarily conducted in non-human primates (see below) and more recently in mice (Keller et al., 2020a; Self et al., 2014; Samonds et al., 2017; Keller et al., 2020b), have elucidated the mechanisms of center-surround modulations in the primary visual cortex (V1). The most frequently observed phenomenon in these studies is suppression, where neuronal responses to stimuli within the center RF are reduced by the presence of specific surrounding stimuli (Knierim and Van Essen, 1992; Levitt and Lund, 1997; Kapadia et al., 1999; Sceniak et al., 1999; Cavanaugh et al., 2002b,c; DeAngelis et al., 1994; Blakemore and Tobin, 1972; Sillito et al., 1995). Suppression is weakest when the peripheral elements oppose the orientation of the central stimulus (Knierim and Van Essen, 1992; Cavanaugh et al., 2002c; Self et al., 2014; DeAngelis et al., 1994), which has been linked, among other things, to the perception of object boundaries (Nothdurft et al., 2000; Lamme, 1995). Surround facilitation, which is less frequently observed, typically occurs when localized, iso-oriented, and collinearly aligned bars are presented in the neuron’s center and surround RF (Levitt and Lund, 1997; Polat et al., 1998; Keller et al., 2020b), and might serve contour integration (Kapadia et al., 1995; Polat et al., 1998; Field et al., 1993).

Contextual modulation of visual responses is influenced by a large array of stimulus features (Angelucci et al., 2017; Nurminen and Angelucci, 2014), including the contrast and the spatial resolution of the stimulus in the center and surround RF (Levitt and Lund, 1997; Kapadia et al., 1999; Sceniak et al., 1999; Polat et al., 1998; Cavanaugh et al., 2002b), the orientation difference between center and surround stimuli (Knierim and Van Essen, 1992; Cavanaugh et al., 2002c), and the spatial resolution of the surround pattern (Li et al., 2006). Although these features interact (e.g. Kapadia et al., 1999), they are often studied independently due to constraints on duration of the experiment. Furthermore, the use of parametric stimuli such as gratings may not optimally drive visual neurons because many neurons in mouse V1 (Walker et al., 2019; Franke et al., 2022; Ustyuzhaninov et al., 2022; Fu et al., 2022) and primate higher visual areas (Pasupathy and Connor, 2001; Bashivan et al., 2019) demonstrate strong selectivity for more complex stimuli, such as corners, checkerboards, or textures.

The strong dependence of contextual modulation on different stimulus features, coupled with the neurons’ preference for complex visual stimuli, underscores the need for an approach to characterize center-surround interactions that does not make strong assumptions on neuronal selectivity and uses ecologically relevant stimuli. Historically, the high dimensionality of natural stimuli and the complexity of interpreting neuronal responses to these stimuli have posed significant challenges. Here, we overcome these challenges by employing a recently developed modeling framework (Walker et al., 2019) and systematically study center-surround modulations in mouse V1 using naturalistic stimuli. This approach involves inception loops –— a closed-loop paradigm that integrates large-scale neuronal recordings, convolutional neural network (CNN) models capable of accurately predicting responses to diverse natural stimuli, *in silico* optimization of non-parametric center and surround images, and *in vivo* verification (Walker et al., 2019; Franke et al., 2022; Bashivan et al., 2019).

Using a data-driven CNN model trained on stimulus-response pairs of experimentally recorded neurons, we synthesized non-parametric surround images that effectively modulated the activity of mouse V1 neurons in response to their preferred stimuli in the center RF, with subsequent *in vivo* verification. Notably, excitatory surrounds *completed* the spatial pattern of the center stimulus, resembling the spatial correlation and congruence of natural scenes (Geisler et al., 2001; Sigman et al., 2001), while inhibitory surrounds *disrupted* the central pattern. We quantified this by using a generative diffusion model to extrapolate natural image statistics from a neuron’s preferred stimulus in the center to the surround, achieving high representational similarity with the model-optimized excitatory surrounds. We additionally tested our approach on macaque V1 by training a CNN model on macaque V1 responses to natural images (Cadena et al., 2023), then applying our synthesis method as well as traditional paradigms using grating stimuli. The synthesized non-parametric surrounds contained complex spatial structures, with completing patterns being more frequent in excitatory than inhibitory surrounds, as observed in mice. Importantly, *in-vivo* experiments in mouse V1 and *in-silico* analyses in macaque V1, respectively, using parametric stimuli replicated previously established center-surround effects with grating stimuli.

Furthermore, to potentially explain the mechanistic basis for excitatory surround pattern completion we demonstrated the presence of “like-to-like” anatomical connections among neurons with minimal RF overlap, employing the “MI-CrONS” functional connectomics dataset (MICrONS Consortium et al., 2021). Finally, we showed that surround excitation and inhibition, driven by pattern completion and disruption, respectively, result as a natural consequence of performing perception as Bayesian inference within a statistical generative model that interprets the stimulus as global objects comprised of local features, thus offering a normative account of the newly discovered center-surround effects.

## Results

### Deep neural network model accurately predicts center-surround modulation of visual responses in mouse primary visual cortex

We combined large-scale population imaging and neural predictive modeling to systematically characterize contextual modulation in mouse primary visual cortex (V1). The experimental and modeling setup was adapted based on (Walker et al., 2019). Specifically, we used two-photon imag-ing to record the population calcium activity in L2/3 of V1 (630×630 µm, 10 planes, 7.97 volumes/s) in awake, head-fixed mice positioned on a treadmill, while presenting the animal with natural images (Fig. 1a,b). For each functional recording, the center RF across all recorded neurons – estimated using a sparse noise stimulus (Jones and Palmer, 1987) – was centered on the monitor (Fig. 1c). This ensured that the center RF of the majority of neurons was within the central area of the monitor. To investigate center-surround interactions in V1 neurons, we presented two types of visual stimuli: full-field natural images (70 x 124 degrees visual angle) and masked natural images (48 degrees in diameter). While the full-field images stimulated both the center (i.e. classical RF) and the surround RF (i.e. the extra-classical surround) of the neurons, the masked images primarily activated the center RF, with a smaller contribution of the surround. Please note that the masked images were not designed to activate solely the center of each neuron without influence from the surrounding area. Instead, using both types of images allowed us to vary the activation levels between the center and surround components of the RF, thus facilitating the learning of surround effects by the model. We used the recorded neuronal activity in response to full-field and masked natural images to train a convolutional neural network (CNN) model to predict neuronal responses as a function of visual input. The model also incorporated eye movements and the modulatory gain effect of the animal’s behavior on neuronal responses (Niell and Stryker, 2010) by using the recorded pupil and running speed traces as input to a shifter and modulator network (Fig. 1d; Walker et al., 2019). A model trained on an example recording session (architecture shown in Fig. 1e) with 7,741 neurons and 4,182 trials (i.e. images) yielded a noise-corrected correlation between model predictions and mean observed responses of 0.73 ± 0.20 (mean±standard deviation; Fig. 1f). This is comparable to state-of-the-art models of mouse V1 (Franke et al., 2022; Willeke et al., 2022; Lurz et al., 2021). Importantly, masking half of the training images as described above improved the model’s prediction of contextual modulation (Fig. 1g): The prediction of how neuronal responses differ between a masked natural image and its full-field counterpart significantly increased when using both full-field and masked natural images during model training (for statistics, see figure legend). Together, this shows that our deep neural network approach accurately captures center-surround modulation of visual responses in mouse primary visual cortex, allowing us to study contextual modulation in the setting of complex and naturalistic visual stimuli.

**Fig. 1.**
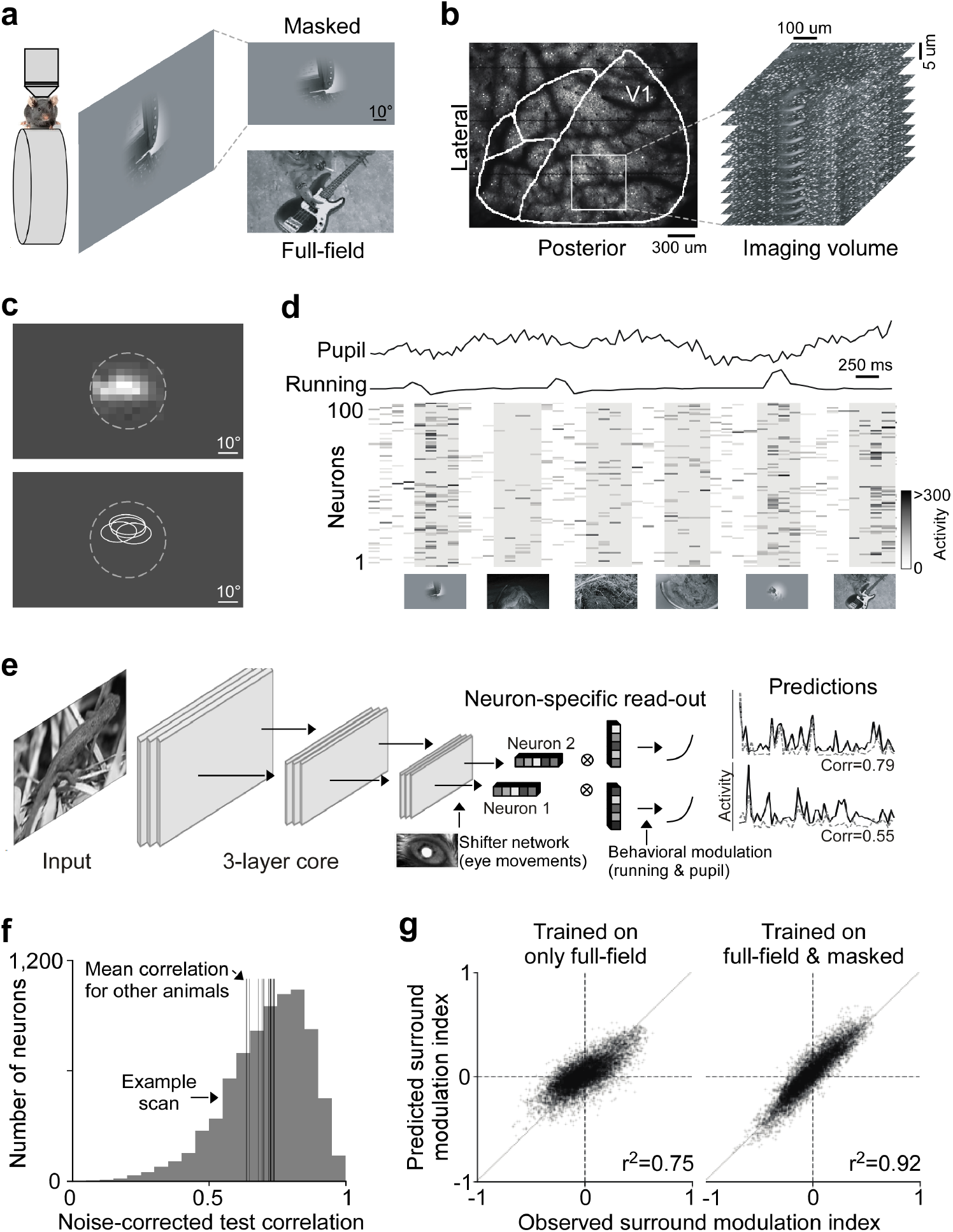
Deep neural network approach captures center-surround modulation of visual responses in mouse primary visual cortex. **a**, Schematic of experimental setup: Awake, head-fixed mice on a treadmill were presented with full-field and masked natural images from the ImageNet database, while recording the population calcium activity in V1 using two-photon imaging. **b**, Example recording field. GCaMP6s expression through cranial window, with the borders of different visual areas indicated in white. Area borders were identified based on the gradient in the retinotopy (Garrett et al., 2014). The recording site was chosen to be in the center of V1, mostly activated by the center region of the monitor. The right depicts a stack of imaging fields across V1 depths (10 fields, 5µstep in z, 630×630µ, 7.97 volumes/s). **c**, Top shows heat map of aggregated population RF of one experiment, obtained using a sparse noise stimulus. The dotted line indicates the aperture of masked natural images. The bottom shows RF contour plots of n=4 experiments and mice. **d**, Raster plot of neuronal responses of 100 example cells to natural images across 6 trials. Trial condition (full-field vs. masked) indicated below each trial. Each image was presented for 0.5s, indicated by the shaded blocks. **e**, Schematic of model architecture. The network consists of a convolutional core, a readout, a shifter network accounting for eye movements by predicting a gaze shift, and a modulator predicting a gain attributed to behavior state of the animal. Model performance was evaluated by comparing predicted responses to a held-out test set to observed responses. **f**, Distribution of normalized correlation between predicted and observed responses averaged over repeats (maximal predictable variability) for an example model trained on data from n=7,741 neurons and n=4,182 trials. Vertical lines indicate mean performance of other animals. **g**, Accuracy of model predictions of surround modulation for only full-field versus full-field and masked natural images. Each test image was presented in both full-field and masked, allowing us to compute a surround modulation index per image per neuron. The modulation indices across images were averaged per neuron. Left and right shows predicted vs. observed surround modulation indices for a model trained on only full-field images and full-field and masked images, respectively. The model trained on both full-field and cropped images predicted surround modulation significantly better than the model trained on only full-field images (p-value<0.001). The total number of training images was the same, and the data was collected from the same animal in the same session.

### CNN model identifies non-parametric excitatory and inhibitory surround images of mouse V1 neurons

We used the trained CNN model as a functional “digital twin” of the mouse visual cortex to identify non-parametric surround images that greatly modulate neuronal activity. For that, we focused on the most ‘exciting’ and most ‘inhibiting’ surround image, which enhances and reduces the response of a single neuron to its optimal stimulus in the center, respectively. The rationale behind this approach was to identify surround images that maximally modulate the encoding of the neuron’s preferred visual feature in the center RF. Please note that the terms ‘excitatory’ and ‘inhibitory’ used to describe optimized surround images do not imply specific synaptic mechanisms but rather describe the functional impact on neuronal activity to the optimal center stimulus. To identify the optimal center stimulus per neuron, we first optimized the most exciting input (MEI) using gradient ascent as previously described (Walker et al., 2019; Franke et al., 2022), corresponding to the non-linear center RF of the neuron. This non-parametric approach of identifying the optimal center stimulus was required because most mouse V1 neurons are not well described by Gabor filters (Fu et al., 2022; Walker et al., 2019). The MEI was optimized using a root mean square (RMS) contrast budget that minimized clipping of pixel values outside the 8-bit range, resulting in an RMS contrast of 12.15±1.35 in 8-bit input space (0 to 255 pixel values). In comparison, the natural images presented during experiments had an RMS contrast of 45.12±17.78. We then used the MEI to define the center RF and consider all visual space beyond the MEI as RF surround (see below for a more detailed discussion on this choice).

To generate excitatory and inhibitory surround images, we held the MEI in the center fixed and only optimized pixels in the surround, starting from Gaussian noise (Fig. 2a). We delineated the borders of the MEI by constructing an MEI mask with smoothed edges. This was achieved through a process of thresholding the MEI at 1.5 standard deviations above the mean, ensuring that the majority of the full-field RMS contrast was encapsulated within the defined mask. During optimization, the center (i.e. MEI mask) of the surround images remained unchanged while the contrast in the surround was redistributed. The RMS contrast budget of the surround was twice the contrast budget we allowed for the MEI, resulting in an RMS contrast of 15 ± 1.5 for the MEI with surround images. This optimization procedure yielded complex features in the RF surround of V1 neurons (Fig. 2b), which were predicted by the model to either enhance or suppress visual responses to optimal stimuli in the center RF (Fig. 2c).

**Fig. 2.**
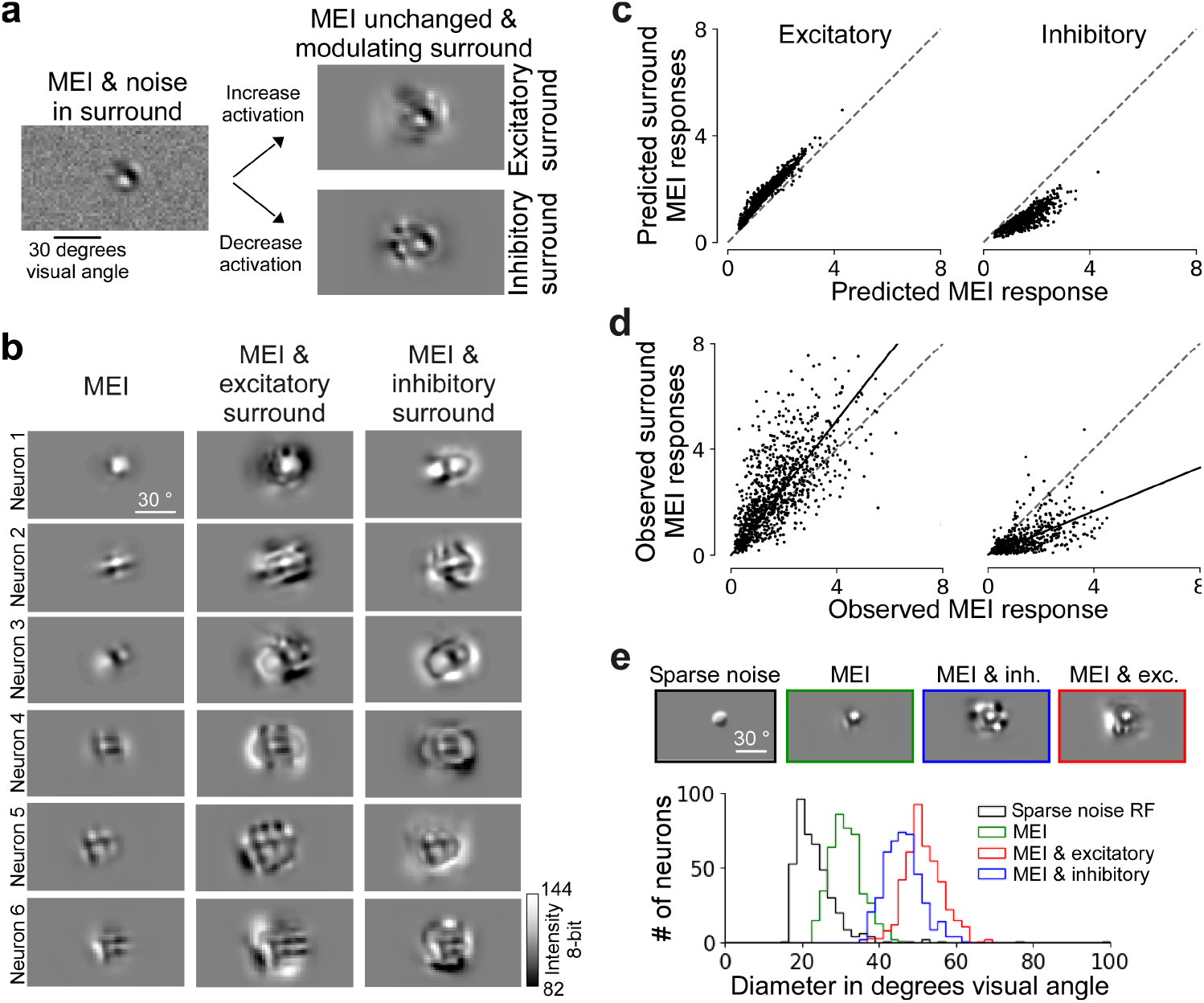
Modeling approach accurately predicts non-parametric excitatory and inhibitory surround images of single neurons in mouse V1. **a**, Schematic of the optimization of surround images. **b**, Panel shows MEI, excitatory surround with MEI, and inhibitory surround with MEI for 6 example neurons. Since the gradient was set to zero during optimization for the area within the MEI mask, the center remained the same as the MEI. **c**, Model predicted responses to MEI and the excitatory (left) and inhibitory (right) surround images (y-axis), compared to the predicted responses to the MEIs (x-axis). Responses are depicted in arbitrary units, corresponding to the output of the model. **d**, Recorded responses to the MEI and excitatory (left) and inhibitory (right) surround (y-axis), compared to the recorded responses to the MEIs (x-axis). For each neuron, responses are normalized by the standard deviation of responses to all images. Across the population, the modulation was significant for both excitatory (n=6 animals, 960 cells, p-value=2.08 × 10^*−*8^, 1.01 × 10^*−*22^, 2.16 × 10^*−*18^, 1.02 × 10^*−*9^, 2.19 × 10^*−*20^, 1.46 × 10^*−*5^, Wilcoxon signed rank test) and inhibitory surround images (n=3 animals, 510 cells, p-value=2.13 × 10^*−*23^, 6.14 × 10^*−*26^, 1.48 × 10^*−*24^). **e**, Diameters of RFs estimated using sparse noise (“center RF”), MEIs, and MEIs with excitatory and inhibitory surround. The means of center RF, MEI, MEI & excitatory, MEI & inhibitory across all neurons are (mean ± s.e.m.): 23.4 degrees ± 0.34 (n=4, 419 cells), 31.3 degrees ± 0.20 (n=4, 434 cells), 51.4 ± 0.23 (n=4, 434 cells), 46.1 ± 0.23 (n=4, 434 cells).

To verify the efficacy of the synthesized surround images *in vivo*, we performed inception loop experiments (Walker et al., 2019; Bashivan et al., 2019): after model training and stimulus optimization, we presented MEIs and the respective surrounds back to the same mouse on the next day while recording from the same neurons, thereby testing whether they effectively modulated neuronal responses as predicted by the model. For a specific recording, we chose 150 neurons from the total population for closed-loop verification. This selection was based on their consistent responses to repeated image presentations and the accuracy of model predictions. We found that the *in silico* predictions (Fig. 2c) matched the *in vivo* results (Fig. 2d, Suppl. Fig. 1): The responses of the neuronal population significantly increased and decreased by the synthesized excitatory and inhibitory surround images, respectively, compared to presenting the MEI alone. The greater variance in the observed responses compared to the predicted ones likely stems from the deterministic nature of our model and the inherent trial-to-trial variability in neuronal responses.

We found that 55.1% of the neurons verified *in vivo* during inception loop experiments were significantly inhibited by their inhibitory surround images across stimulus repetitions. In contrast, only 28.4% of neurons were significantly facilitated by their excitatory surround images. Critically, less than 3% of neurons were significantly modulated in the direction opposite to what the model predicted. We also performed a subset of experiments using a higher contrast budget for center and surround MEIs (22.23±3.38 for MEIs, 26.86±3.65 and 29.22±4.26 for MEI with excitatory and inhibitory surround, respectively), while keeping the ratio between center and surround contrast unchanged (Suppl. Fig. 2a). While the strength of surround modulation decreased for higher contrast levels, excitatory and inhibitory surround MEIs still significantly modulated center responses of mouse V1 neurons (Suppl. Fig. 2b). Together, these results from the inception loop experiments demonstrate the accuracy of our CNN model in synthesizing effective non-parametric modulatory surround images of mouse V1 neurons.

To investigate the ecological relevance of the center-surround modulation observed with non-parametric images, we examined if similar modulation occurs in mouse V1 neuronal activity with natural images (Suppl. Fig. 3). We specifically targeted natural images that mimic the neuron’s preferred center feature, akin to the optimized surrounds. For each neuron, we screened 5,000 natural images masked with the neuron’s MEI mask, selecting those that elicited strong activation (>80% relative to the maximum excitatory input or MEI; Suppl. Fig. 3a). After replacing the center of these images with the MEI and adjusting them to match the average size and contrast of the excitatory and inhibitory surrounds, we evaluated the modulation strength by presenting these modified natural images to both the model and the animal. We then measured the *in-silico* and *in-vivo* responses, comparing them to the responses elicited by the MEI alone. Our findings revealed that certain natural surrounds can either enhance or reduce V1 responses to the preferred visual feature, paralleling the effects seen with synthesized surrounds (Suppl. Fig. 3b,c). Generally, the modulation strength elicited by the synthesized images exceeded that of the natural surrounds. These results strongly indicate that our model-derived surrounds are ecologically relevant, effectively mimicking the modulation of V1 responses by natural image surrounds.

We performed a number of control experiments to verify that the observed response modulations indeed originated from activating the surround of the neurons. As described above, we used the MEI as an approximation of the center RF and defined visual space beyond the MEI as surround RF. To relate our definition of center RF and surround to definitions used previously, we first demonstrated that the synthesized surround images indeed extend beyond the center RF of the neurons (Fig. 2e), identified using a well-established stimulus for RF mapping. Specifically, we estimated each neuron’s center RF as the minimal response field (MRF) using a sparse noise stimulus (Jones and Palmer, 1987) and compared its size to the size of the MEI and the excitatory and inhibitory surround, respectively. The MRF was, on average, smaller than the MEI, suggesting that the MEI itself corresponds to an overestimation of the center RF. Lowering the contrast of the sparse noise stimulus to more closely match the contrast of the MEI did not change the distribution of MRF sizes (Suppl. Fig. 2c). Importantly, both the excitatory and inhibitory surround were much larger than the MRF, indicating that the modulatory effect on neuronal activity we observed by the surround images was indeed elicited by activating the surround component of V1 RFs.

In line with this, in additional control experiments we showed that the response modulation did not solely originate from the region directly adjacent to the MEI but further increased both *in silico* and *in vivo* when considering the full surround region (Suppl. Fig. 4). Finally, we showed that increasing the contrast in the center was more effective in driving the neurons than adding the same amount of contrast in the surround of the image (Suppl. Fig. 5), consistent with the idea that the enhancement in neuronal response from the surround is weaker than from the center (Allman et al., 1985; Cavanaugh et al., 2002a; Jones et al., 2001; Knierim and Van Essen, 1992). Together, these results demonstrate that the observed response modulation by model-derived surround images originates from activating the surround RF of V1 neurons.

### Pattern completion and disruption shaped by natural image statistics characterize excitatory and inhibitory surround images

Center-surround modulation of visual activity corresponds to a neuronal implementation for integrating visual information across space, thereby providing context for visual processing. So far, models of contextual modulation have largely focused on parametric stimuli for visual cortex, such as gratings (but see e.g. Coen-Cagli et al., 2012), perhaps due to the lack of tools that allow unbiased and systematic testing of such high-dimensional visual inputs. Here, we used our data-driven model and the optimized surround images to systematically investigate the rules that determine contextual excitation versus inhibition in a naturalistic setting.

We observed that excitatory surround images demonstrated greater congruence with the MEI in the center compared to inhibitory surround images (Fig. 3a). Specifically, the spatial patterns within the MEI, such as orientation (e.g., neurons 2 and 3), were predominantly preserved by the excitatory surround, whereas the inhibitory surround tended to disrupt these patterns. This pattern completion and disruption was also evident for more complex spatial structures like grid patterns (neuron 1), which were completed by the excitatory surround and fragmented by the inhibitory surround. Notably, the congruent patterns observed for MEIs with excitatory surrounds echo the well-documented phenomenon wherein natural images frequently exhibit congruent structures that delineate object contours and create continuous patterns (Geisler et al., 2001; Sigman et al., 2001). Consequently, we propose the hypothesis that MEIs accompanied by excitatory surrounds may share statistical characteristics with, and appear perceptually similar to, natural images—– more so than MEIs with inhibitory surrounds. More broadly, we suggest that the rules that govern surround excitation and inhibition may be described as completion and disruption of spatial patterns according to natural image statistics.

**Fig. 3.**
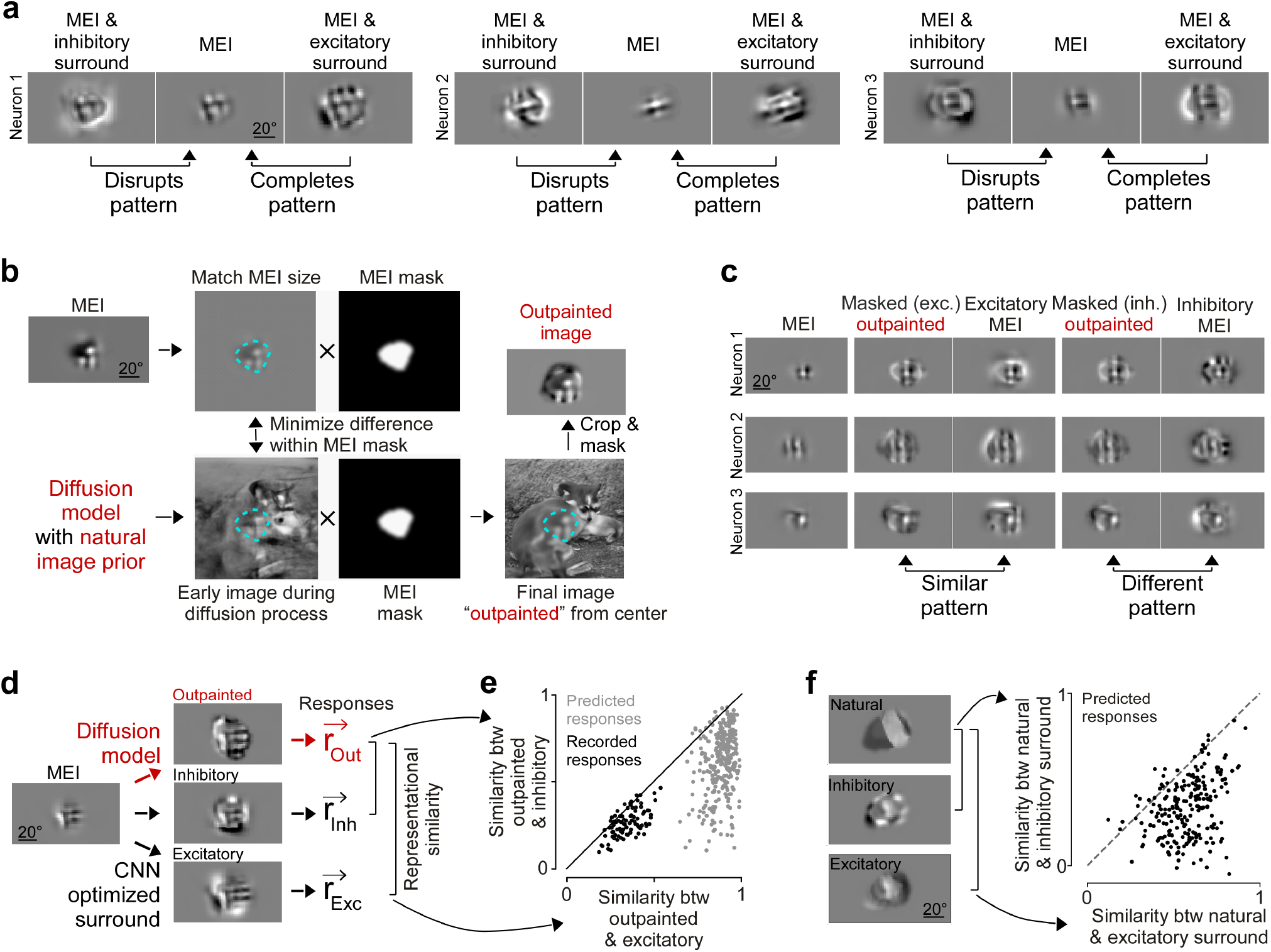
Completion and disruption of natural image statistics characterize excitatory and inhibitory surround images. **a**, MEI with excitatory and inhibitory surround of four example neurons, illustrating that excitatory and inhibitory surround images complete and disrupt, respectively, spatial patterns of the MEI. **b**, Schematic illustrating how we used a diffusion model with a natural image prior to outpaint spatial patterns of the MEI into the surround. The diffusion process included an additional loss function, which minimized the difference between MEI and the generated image within the MEI mask (L2 norm). This resulted in a final image outpainted from the center, which includes MEI features in the center and naturalistic features in the surround. The outpainted surround image was created independent on the neuron’s activation and instead maximized the consistency between center and surround with respect to natural image statistics. **c**, MEI, optimized excitatory and inhibitory surrounds, and outpainted surrounds, masked either using the excitatory or inhibitory surround mask, for three example neurons. **d**, Schematic illustrating how we estimated representational similarity of MEI and surround images. We presented MEIs, optimized and outpainted surround images to the animal in closed loop experiments and obtained a population response for each presented image (r). We then estimated representational similarity between outpainted and optimized surround images by estimating the Cosyne similarity between image pairs. **e**, Representational similarity (as Pearson’s correlation coefficient between neuronal responses) of outpainted surround images to excitatory and inhibitory surround. The black dots indicate data from *in vivo* closed-loop experiments (n=1 animal, 90 cells, p-value=4.20 × 10^*−*4^, Wilcoxon signed rank test). The gray dots indicate data from *in silico* experiments (n=2 animals, 300 cells, p-value=3.04 × 10^*−*5^). **f**, Representational similarity (as Pearson’s correlation coefficient between neuronal responses) of natural image surround to excitatory and inhibitory surround. One exciting natural image and surround of one example neuron (left) and representational similarity (as Pearson’s correlation coefficient, right; p-value=5.28 × 10^*−*35^, two-sided Wilcoxon signed rank test, n=3 animals, 219 neurons) of natural surround images with excitatory and inhibitory surround. Each dot represents the mean across natural surrounds per neuron.

To evaluate our hypothesis, we extrapolated the spatial patterns of the MEI from the center into the surround using a generative diffusion model (Pierzchlewicz et al., 2023) trained on a dataset of natural images ((Fig. 3b); Dhariwal and Nichol, 2021). This process, known as “outpainting” in computer vision, generated surround images based on the statistical properties of natural images. It is crucial to note that these outpainted surrounds solely relied on the statistics of natural images learned by the diffusion model. In particular, they are independent of the CNN model employed to predict neuronal activity and optimize MEIs. This ensures that testing our hypothesis was not influenced by the predictive model itself. For each neuron, we started with the MEI in the center and outpainted the surround 40 times, resulting in 40 unique outpainted surround images per neuron through the diffusion model’s stochastic sampling. We then masked the outpainted images using the surround MEI masks and adjusted the contrast, such that the outpainted surround images had the same size and contrast as the MEI with excitatory and inhibitory surround, respectively.

We found that the outpainted surround images, averaged across the 40 unique images per neuron, qualitatively looked more similar to the excitatory than the inhibitory surrounds (Fig. 3c), in line with our hypothesis stated above. To quantify the similarity of CNN-optimized and outpainted surrounds, we computed the “representational similarity” (Kriegeskorte et al., 2008) for a given pair of images in the V1 neuronal response space. We chose to use representational similarity instead of pixel-wise correlation to quantify similarity between images because (i) the representational space more closely mimics similarity at the representational level of interest (mouse V1) and (ii) this process removes image features that are irrelevant to the visual system, such as high spatial frequency noise. We performed closed-loop experiments and presented the outpainted surround images back to the animal, in addition to the MEIs with excitatory and inhibitory surrounds as described above. For each presented image, we obtained a vector of recorded neuronal responses, averaged across repeated trials and computed the cosine similarity between the mean response vectors of an image pair, i.e. we correlated the population response vectors of outpainted and excitatory surround or the population response vectors of outpainted and inhibitory surround (Fig. 3d). We found that the outpainted surround images exhibited a high representational similarity to the MEI with excitatory surrounds, while the similarity to the MEI with inhibitory surrounds was much weaker (Fig. 3e). This trend was even more pronounced when using the CNN-model predicted responses instead of the recorded responses for estimating the representational similarity between outpainted and excitatory and inhibitory surround images (Fig. 3c). Please note that the representational similarity metrics derived from predicted responses of outpainted and inhibitory surrounds exhibited considerable variability across images. Nonetheless, there was a significant correlation between the representational similarities derived from predicted and recorded responses. The outpainted images displayed central spatial structures that resemble, but are not identical to, the MEI, and given the model’s sensitivity to variations in its predicted MEIs, this could account for the observed variability in predicted responses, while the recorded neuronal population may remain invariant to these minor modifications.

To further demonstrate that MEIs with excitatory surrounds were indeed more closely aligned with natural images than those with inhibitory surrounds, we employed the representational similarity metric introduced earlier on natural images directly. For each neuron, we began by identifying highly activating natural image crops located in the center RF. We then extended these images into the surround using the masks from the CNN-optimized surround images, and adjusted the contrast to align with the optimized center-surround images. We then presented both optimized and natural images to the model. In line with our predictions, this analysis showed that natural surround images featuring the neuron’s preferred center exhibited greater similarity to MEIs with excitatory surrounds than to those with inhibitory surrounds (Fig. 3f).

To check whether excitatory and inhibitory surrounds are also characterized by first order image statistics like mean luminance, in addition to pattern completion and disruption, we compared the distribution of pixel values of excitatory and inhibitory surround MEIs. This revealed that the pixel value distributions of excitatory and inhibitory surround MEIs did not significantly differ from one another, suggesting that negative contrasts or not generally more exciting than positive contrasts. In addition, the mean pixel value of the excitatory surround MEI was negatively correlated with the mean pixel value of the inhibitory surround MEI, indicating that excitatory and inhibitory surround MEIs have opposite mean luminance. For example, if the excitatory surround MEI of a neuron is dominated by negative contrast, then the inhibitory surround MEI is dominated by positive contrast, and vice versa. This analyses on first order statistics complements our analysis above on higher order natural image statistics.

Taken together, our results demonstrate that surround excitation and inhibition in mouse primary visual cortex can be characterized by pattern completion and disruption, respectively, based on natural image statistics. This yields a novel relationship between natural image statistics and modulation of neuronal activity in the visual system.

### Iso- and ortho-oriented surround grating stimuli generally suppress center drifting grating responses

Our findings challenge the prevailing view that congruent spatial patterns, such as a surround grating matching the orientation of the center grating, are generally more inhibitory than orthogonal spatial patterns (e.g. DeAngelis et al., 1994; Cavanaugh et al., 2002c; Self et al., 2014), and that excitatory effects of the surround are rare (reviewed in Angelucci et al., 2017).

However, it is important to note that surround effects appear to be influenced by numerous factors, including the contrast of the center and surround stimuli, the type of stimulus used, and others (reviewed in Angelucci et al., 2017). For example, unlike our study, which uses natural images and a non-parametric approach to identify modulating surround images, most research on contextual modulation in the visual cortex has relied on well-defined parametric stimuli, such as drifting gratings. To investigate the extent to which the discrepancies between our results and previous studies could be attributed to differences in the types of visual stimuli used, we conducted experiments presenting both sinusoidal drifting gratings and natural images to the same set of mouse V1 neurons. Specifically, we presented drifting gratings with predetermined spatial and temporal frequencies (0.05 cpd and 1.2 Hz, based on Self et al. (2014)) at the screen’s center (diameter 20 degrees visual angle), either in isolation or accompanied by iso- or orthogonal-oriented gratings in the surround, maintaining the same frequencies as the center stimulus (Fig. 4a). Concurrently, we presented both masked and unmasked natural images to the same neurons, as detailed above. This approach enabled a direct comparison between the effects of surround modulation induced by drifting gratings and natural images. In our analysis, we focused exclusively on neurons with RFs centered on the screen, directly overlapping with the stimulus area of the center drifting grating. Our findings indicate that both iso- and orthogonal-oriented surround gratings generally suppress the neuronal response to central drifting gratings, although there was considerable variability across individual neurons (Fig. 4b). In addition, iso-oriented gratings were on average more effective in suppressing center responses than orthogonal oriented gratings. Crucially, for anatomically matched neurons across stimuli, our model trained on responses to natural images predicted the excitatory and inhibitory surround modulation patterns described above (Fig. 4c). Our results using grating stimuli thus align with previous research, suggesting an interplay between stimulus statistics and neuronal response modulation. This underscores the importance of employing a variety of visual stimuli to fully understand the dynamics of contextual modulation in the visual cortex.

**Fig. 4.**
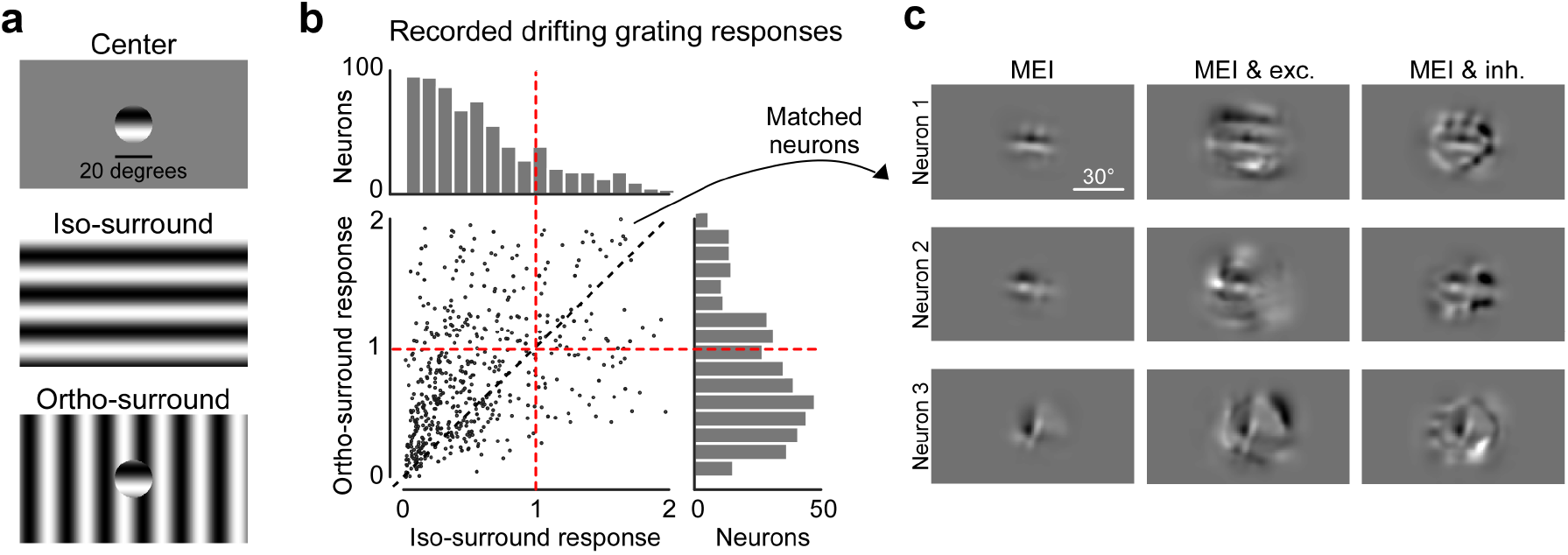
Surround suppression in mouse V1 using drifting grating stimuli. **a**, Schematic illustrating visual stimuli used in the grating experiment, including center-only, center with iso-oriented surround and center with orthogonal surround. **b**, Scatter plot of surround modulation by iso-oriented and ortho-oriented surround. For each neuron, the responses to center-surround stimuli are normalized by responses to the center stimulus alone. Both iso-oriented and orthogonal surround are more suppressive, with the iso-oriented surround eliciting lower neuronal responses (Wilcoxon signed rank test, p-value=3.22 × 10^*−*50^).

### Predictive model trained on macaque V1 responses to natural images reproduces center-surround interactions discovered in mice

Next, we investigated whether the center-surround effects observed in mouse primary visual cortex are also present in macaque visual cortex, where much of the previous research has been conducted. We used an existing dataset of macaque V1 single neuron spiking activity to natural images (n=458 neurons, n=2 macaques, Cadena et al. (2023)) and trained a CNN model to predict spiking activity in response to these images (Fig 5a,b). Our model achieved a mean correlation of 0.74 with trial-averaged experimentally recorded responses (Fig 5c), slightly outperforming existing models for macaque V1 (Cadena et al., 2023). We focused further analysis on the best-predicted neurons, those exceeding an inclusion threshold of 0.75 correlation between predicted and trial-averaged measured activity (n=252 neurons). Similar to our approach with mice, we regarded the model as a functional “digital twin” of macaque V1 and employed it for detailed in-silico analysis of contextual modulation of visual responses. It is important to note that all further analyses are performed in the model, and not directly in experiments with the animals.

**Fig. 5.**
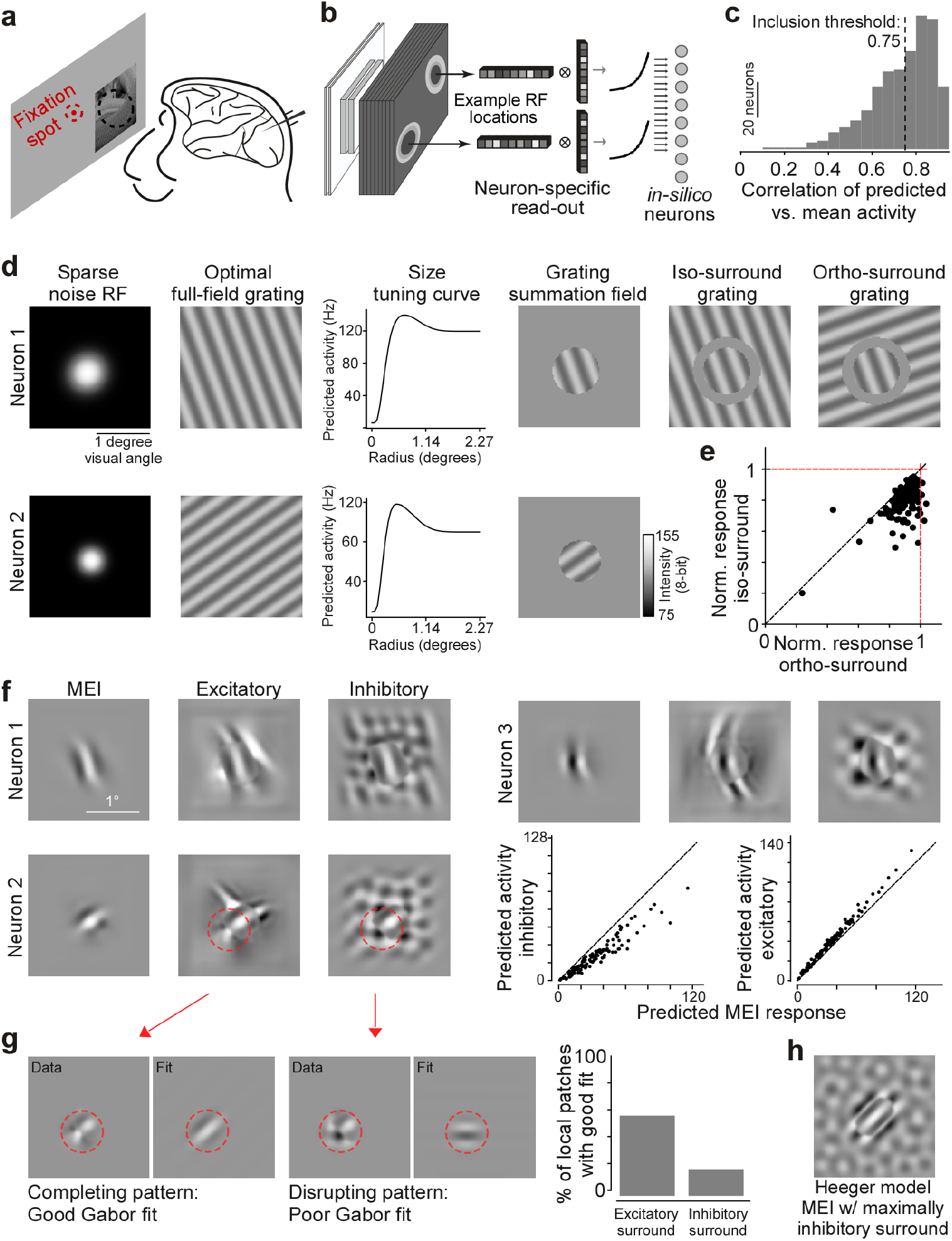
Predictive model trained on macaque V1 responses to natural images reproduces center-surround interactions discovered in mice. **a**, Schematic of experimental setup: awake, head-fixed macaques were presented with grey-scale natural images from the ImageNet database at a parafoveal eccentricity while focusing on a fixation spot. Neuronal spiking activity was recorded using linear probes. Data from (Cadena et al., 2023). **b**, Schematic of model architecture. A ConvNext CNN model was trained on the collected experimental data to predict the spike rate of the recorded neurons to natural images. **c**, Histogram of correlation of model predictions to trial averaged responses of held out test-dataset. Only neurons with correlation above the inclusion threshold of 0.75 are considered for the subsequent *in-silico* experiments. **d** Results of classical experiments performed in-silico for two neurons. From left to right: Gaussian fitted sparse noise RF, optimal full-field grating, size tuning curve, grating summation field (GSF) and GSF in the center, with iso- and ortho-oriented surround gratings (orientation contrast). **e**, Scatter plot summarizing results of orientation contrast experiment. Plot shows model predicted responses to GSF with ortho- and iso-oriented surround gratings, normalized per neuron based on the firing rate to the GSF alone. **f**, Optimized MEI and excitatory and inhibitory surround stimuli with MEI for 3 example neurons. Scatter plots show model predicted responses to MEI versus to MEI with surround images. **g**, Images on the left illustrate quantification of pattern completion and disruption for excitatory and inhibitory surround stimuli with MEI: local patches (data) at the border of center and surround are extracted from the optimized stimuli and then fitted with a Gabor (fit). The right panel displays the percentage or local patches with good Gabor fit. A 0.3 MSE threshold was chosen to discriminate between good Gabor fits (corresponding to MEI pattern continuation in surround) and poor Gabor fits (corresponding to MEI pattern disruption in surround). **h** MEI with maximally inhibitory surround of example neuron obtained from a simulated dataset and a Heeger model of divisive normalization (Heeger, 1992). We simulated 10,000 linear-non-linear simple cells as Gabor filters, with randomly sampled position and orientation.

We validated our model’s accuracy in capturing well-known center-surround interactions in macaque V1 through a series of experiments with established parametric grating stimuli (Fig 5d). We mapped each neuron’s RF using a sparse noise stimulus and identified its preferred spatial frequency and orientation via presentation of full-field sinusoidal gratings (gratings spanned pixel values between 102 and 127 in 8-bit range). We then conducted a size tuning experiment, presenting gratings of the preferred orientation and spatial frequency to each selected neuron, masked by a disk of increasing radius centered on the sparse noise RF. This revealed that most neurons exhibited surround suppression, where their response initially increased as the radius of the grating expanded, then decreased again. We defined the grating summation field (GSF) as the smallest grating that elicited 95% of the maximum activation (Cavanaugh et al., 2002c), which was typically larger (mean across neurons 0.96 degrees) than the sparse noise estimated RFs (mean across neurons 0.61 degrees; Fig 5d). Further, we performed an orientation contrast experiment, presenting each selected neuron with stimuli composed of its GSF paired with either an iso-oriented or ortho-oriented grating in the surround, separated from the center by a moat of 0.23 degrees visual angle (Fig 5d, top right). This experiment demonstrated, on average, stronger suppression for iso-oriented than for ortho-oriented surrounds, and revealed surround facilitation for ortho-oriented surrounds in a small subset of neurons. These findings align with previous research conducted in macaque V1 (reviewed in Angelucci et al., 2017), confirming that our CNN model accurately learns and reproduces classic experiments on center-surround interactions.

We next focused on identifying non-parametric surround images that optimally inhibit and excite, respectively, each neuron’s firing to its preferred visual feature in the center RF, following the same approach to that previously detailed in our mouse experiments. First, we identified each neuron’s MEI using optimization in pixel space, with a pixel standard deviation approximately matching the standard deviation of the gratings used in the above experiments. Consistent with earlier findings (Fu et al., 2022), the majority of macaque V1 MEIs resembled Gabor patterns (Fig 5f, Suppl. Fig. 6). The size of the MEIs (mean across neurons 0.88 degrees) was larger than the sparse noise RF and comparable to that of the GSF, suggesting that the MEI provides a reliable approximation of the neuron’s center RF extent. Subsequently, we synthesized modulatory surround images by keeping the MEI fixed and optimizing only the surrounding pixels to either increase (excitatory) or decrease (inhibitory) the neuron’s response to the MEI in the center (Fig 5f,, Suppl. Fig. 6). The excitatory surround patterns typically continued the Gabor pattern present in the center (e.g. neurons 1 and 3), particularly along the axis of the preferred orientation in the center. For many neurons, the excitatory surround additionally included local Gabor-like features with a different orientation along the flanking sides of the MEI. Those were often orthogonal with respect to the MEI orientation, thereby adding complexity to the pattern (e.g. neuron 2). Conversely, inhibitory surrounds often displayed a texture-like grid pattern that tended to disrupt the central Gabor pattern. We quantified pattern completion and disruption for excitatory and inhibitory surrounds by taking advantage of the fact that most individual macaque V1 neurons’ MEIs are well-described by a Gabor filter. Specifically, we extracted local patches from the optimized images at the border between MEI and the surround and fitted these patterns with Gabor functions (Fig 5g left panel). The reasoning behind this analysis is as follows: if we can fit the local patch at the border between MEI and surround with a Gabor function with little error, this indicates pattern continuation in the near surround adjacent to the MEI. Conversely, if Gabor patterns present in the MEI do not continue in the near surround, the Gabor fit on the respective local patch will be poor, suggesting pattern disruption. Our results indicate that such local Gabor pattern continuation occurs much more frequently for the excitatory surround than for the inhibitory one (see Fig 5g, right and Suppl. Fig. 6).

Previous research has indicated that the inhibitory surround of V1 neurons is influenced by the collective activity of a diverse array of V1 neurons, potentially facilitating divisive normalization—a mechanism that standardizes each neuron’s responses relative to its neighboring activity (Heeger, 1992; Carandini and Heeger, 2011). Accordingly, we hypothesized that the texture-like patterns frequently observed in inhibitory surround images could correspond to stimuli that optimally drive a population of Gabor neurons characterized by varying orientation preferences and spatial positions. To investigate this hypothesis, we constructed a model comprising a population of simple cells represented as linear-non-linear (LN) neurons featuring Gabor-shaped RFs, with random variations in position and orientation. Subsequently, we implemented a simple divisive normalization model (Heeger, 1992) centered on an LN simple cell, wherein the response is divisively normalized by the activity of the neuron population. This process yielded both the MEI and its corresponding maximally inhibitory surround of this LN simple cell (Fig. 5h). The resultant image exhibited pattern disruption and a texture-like appearance reminiscent of the inhibitory surrounds observed in macaque V1 neurons. Our findings lend support to the notion that the patterns evident in most inhibitory surrounds could emerge from the cumulative activity of a population of neurons, potentially serving divisive normalization mechanisms.

These insights from mouse and macaque V1 suggest that within the framework of natural images as visual stimuli and a non-parametric analytical approach, pattern completion and disruption drive surround excitation and inhibition, respectively, in the primary visual cortex of both mice and macaques. That being said, in macaques, there is a great diversity of surround patterns, particularly within excitatory surrounds, including the frequent appearance of orthogonally oriented features. By integrating both established parametric stimuli and innovative non-parametric methods, our findings not only align with but also significantly enhance the existing understanding of surround interactions in V1.

### Circuit-level dissection using the MICrONS dataset identifies “like-to-like” connections across broad spatial scale as potential mechanism of pattern completion

To further understand the mechanisms at the circuit level contributing to the established rules governing contextual modulation in mouse V1, we integrated functional recordings with anatomical analyses. For that, we used the “MICrONS” dataset, which includes responses of over 75,000 neurons to full-field natural movies along with reconstructed sub-cellular connectivity from electron microscopy data (MICrONS Consortium et al., 2021). Crucially, a “dynamic” model–a recurrent neural network (RNN) representing a digital twin of this portion of the mouse visual cortex–demonstrates not only high predictive accuracy for responses to natural movies but also robust out-of-domain performance with other stimulus classes such as drifting Gabor filters, directional pink noise, and random dot kinematograms. This allows for the presentation of novel stimuli to the digital twin model, facilitating a detailed exploration of how specific functional properties correlate with the underlying neuronal connectivity and anatomical characteristics.

Here, we evaluated whether the dynamic model trained on the MICrONS dataset accurately replicates the center-surround effects observed in our experiments, thereby serving as a tool for circuit-level analysis of these interactions. We used the MICrONS dynamic model to simulate responses to both full-field and masked natural images used during our experiments (Fig. 6a), then trained a model on these predicted responses and used it to optimize MEIs and their excitatory and inhibitory surrounds for the neurons (“dynamic to static”). Our findings revealed that the excitatory and inhibitory surround images, respectively, complete and disrupt the spatial patterns present in the MEI, in line with our experimental results (Fig. 6a, right). To ensure that this “dynamic to static” approach indeed generates surround images that accurately modulate neuronal activity as predicted, we conducted additional closed-loop experiments. These experiments are essential for verifying the model’s predictions with new visual stimuli not included in the original training set. We recorded neuronal responses to static natural images and the same natural movies used in the MICrONS dataset. We subsequently trained two CNN models: one directly on the recorded responses to natural images, and another on responses predicted by a dynamic model that had been trained from scratch on recorded responses to natural movies. The MEIs and surround images derived from these two models showed remarkable perceptual similarity (Fig. 6b, left). When these MEIs and their corresponding surround images were presented back to the animal, the “dynamic to static” generated surrounds modulated neuronal activity in expected ways—increasing activity with excitatory surrounds and decreasing it with inhibitory surrounds (Fig. 6b, right). This shows that our results are applicable to natural movie data, thus validating the use of the MICrONS dataset to explore neuronal circuits that underlie center-surround interactions.

**Fig. 6.**
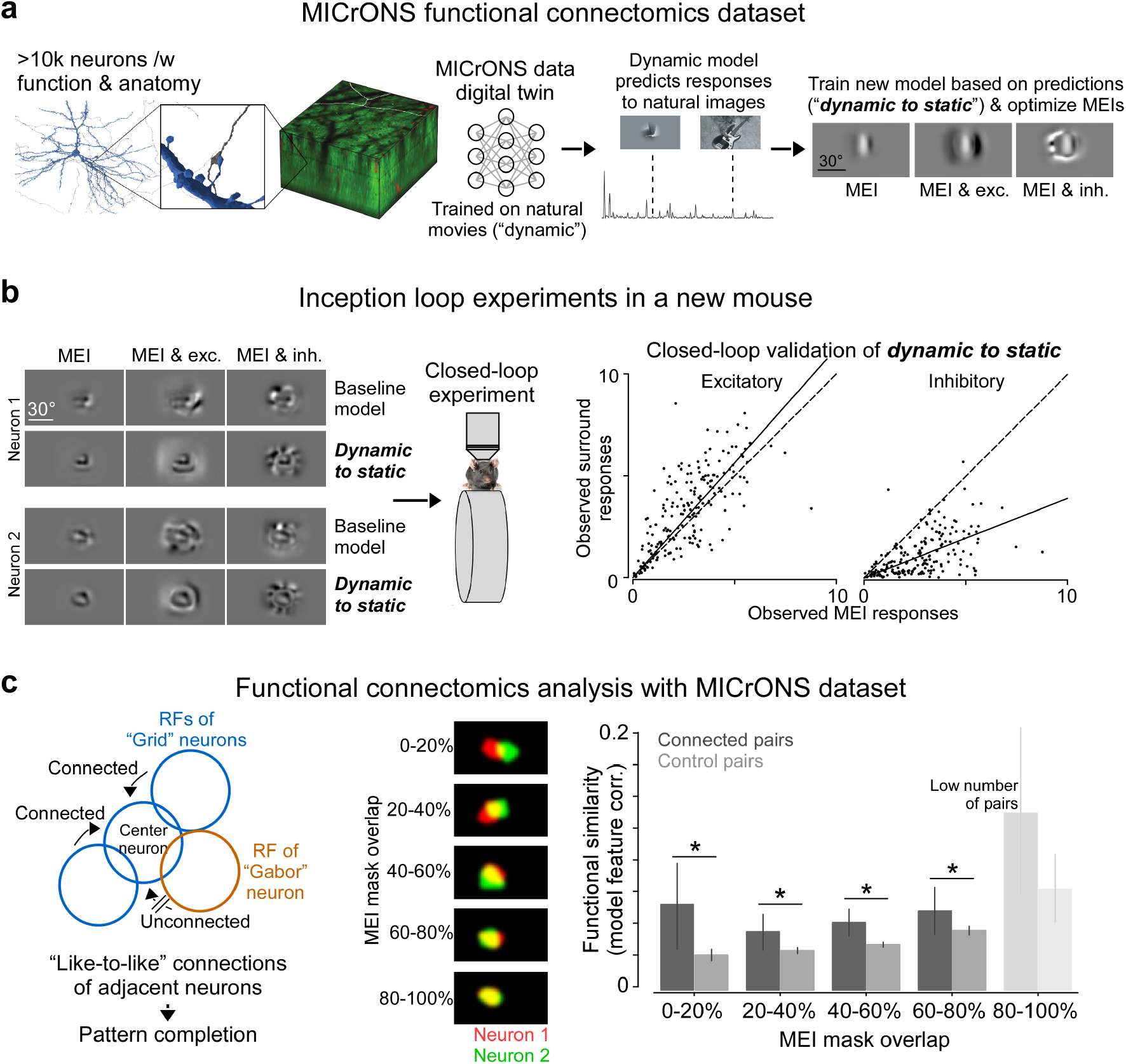
**a**, Schematic shows the MICrONS functional connectomics dataset (MICrONS Consortium et al., 2021), which includes responses of >75k neurons to full-field natural movies and the reconstructed sub-cellular connectivity of the same cells from electron microscopy data. We used the MICrONS digital twin (Wang et al., 2023) trained on natural movies (“dynamic” model) to predict responses to natural images used in our experiments. We then trained a new model based on these predictions (“dynamic to static”) and optimized MEIs and surround images. **b**, Verification of center-surround effects of the MICrONS digital twin (panel (a)). Left shows MEIs and excitatory and inhibitory surround images for two example neurons, optimized using our baseline model used for all mouse experiments and the “dynamic to static” pipeline described in panel (a). Neurons were matched across natural movie and image recordings based on their position in a high-resolution 3D stack. MEIs and surround images were presented to the animal in closed-loop experiments. Right shows observed MEI responses plotted versus observed responses to MEI with excitatory and inhibitory MEI, using the “dynamic to static” method for image synthesis. Surround modulation was significant for both excitatory surround (n=1 animal, 200 cells, p-value=2.12 × 10^*−*9^, Wilcoxon signed rank test) and inhibitory surround (n=1 animal, 200 cells, p-value=8.87 × 10^*−*32^). **c**, Left shows schematic illustrating hypothesis. Anatomical connections between adjacent neurons with high functional similarity (“like-to-like”) could underlie pattern completion for the excitatory surround. To investigate this, we split pairs of neurons in V1 L2/3 with proof-read connectivity from the MICrONS dataset into groups based on the amount of overlap between their MEI masks (middle). We then compared the feature similarity among pairs with different amounts of MEI overlap (right). The significance is derived from Welch’s t-test and p values are corrected for multiple-test correction. Asterisks indicate p-value < 0.05. We used a Poisson generalized linear model to predict number of synapses from mask overlap and feature similarity. This revealed that both mask overlap and feature similarity are significantly larger than zero, while the weight for the interaction between mask overlap and feature similarity is not significantly different from zero.

Prior studies have established that excitatory cortical neurons are more likely to form anatomical connections if they exhibit functional similarities, a phenomenon described as like-to-like connections (Ko et al., 2011; Cossell et al., 2015; Lee et al., 2016; Scholl et al., 2021). Our observation that a completing surround pattern is excitatory suggests that neurons with similar functional characteristics are inclined to connect even when their RFs do not overlap. For instance, a neuron preferring feature A would likely receive excitatory connections from other feature A-preferring neurons in the surrounding area, effectively completing the central pattern (Fig. 6c, left). To evaluate this hypothesis, we analyzed the RF overlap and functional similarity of neuron pairs within the MICrONS dataset, consisting of 624 neurons and 793 synapses. These pairs were either anatomically connected (‘connected’) or randomly pooled from the dataset, irrespective of their connectivity (‘control’). We approximated each neuron’s RF using the MEI and assessed the functional similarity between neuron pairs by measuring the cosine similarity of the neuron-specific feature weights of the dynamic model. In alignment with existing literature, our results demonstrated that connected neuron pairs displayed greater functional similarity compared to control pairs. Furthermore, our analysis revealed the persistence of this effect across a spectrum of RF overlaps. Notably, even neuron pairs with minimal RF overlap (0-20%) exhibited higher functional similarities relative to control pairs. To further quantify these relationships, we applied a generalized linear model to model the synaptic connectivity based on functional similarity and RF overlap, as well as their interaction. We found that functional similarity and RF overlap independently predicted the number of synapses between neuron pairs, since the interaction term between functional similarity and RF overlap did not significantly contribute to predicting synapse numbers. This suggests that functionally similar neurons are more likely to form synapses than control pairs, irrespective of their RF overlap. This observation supports the presence of an excitatory completing surround pattern and offers valuable insights into the circuit-level mechanisms involved in such neuronal interactions.

### Perception as Bayesian inference explains observed center– surround effects

Finally, we linked our observed center-surround effects to normative, first-principles theories of perceptual inference — specifically posterior inference, where the brain updates its internal beliefs (posterior) in light of new sensory input (evidence) against prior beliefs or experiences (prior probabilities). The primary goal of perception is to infer useful features from the environment. Due to the inherent ambiguity and noise in sensory stimuli it is advantageous to integrate the information from sensory stimuli with pre-existing knowledge or beliefs about the environment (Von Helmholtz, 1867). A principled way to accomplish this integration is through Bayesian inference on relevant world variables (latent features) that are part of a statistical generative model of the world (Knill and Richards, 1996; Kersten et al., 2004; Lee and Mumford, 2003; Fiser et al., 2010). The theory does not claim that the brain maintains a generative model itself, but that neuronal activity represents the result of the process of “inverting” a generative model, that is, inferring possible world configuration that could have led to the transmitted sensory signals from, e.g., the retina. In statistical terms, this can be formalized by computing a posterior over the world variables given the sensory evidence. Here, we show that surround excitation and inhibition elicited by completing and disrupting surround patterns respectively, are a natural consequence of performing Bayesian inference in a generative model of the stimulus that explains the stimulus as global objects consisting of local features (Fig 7a-c).

**Fig. 7.**
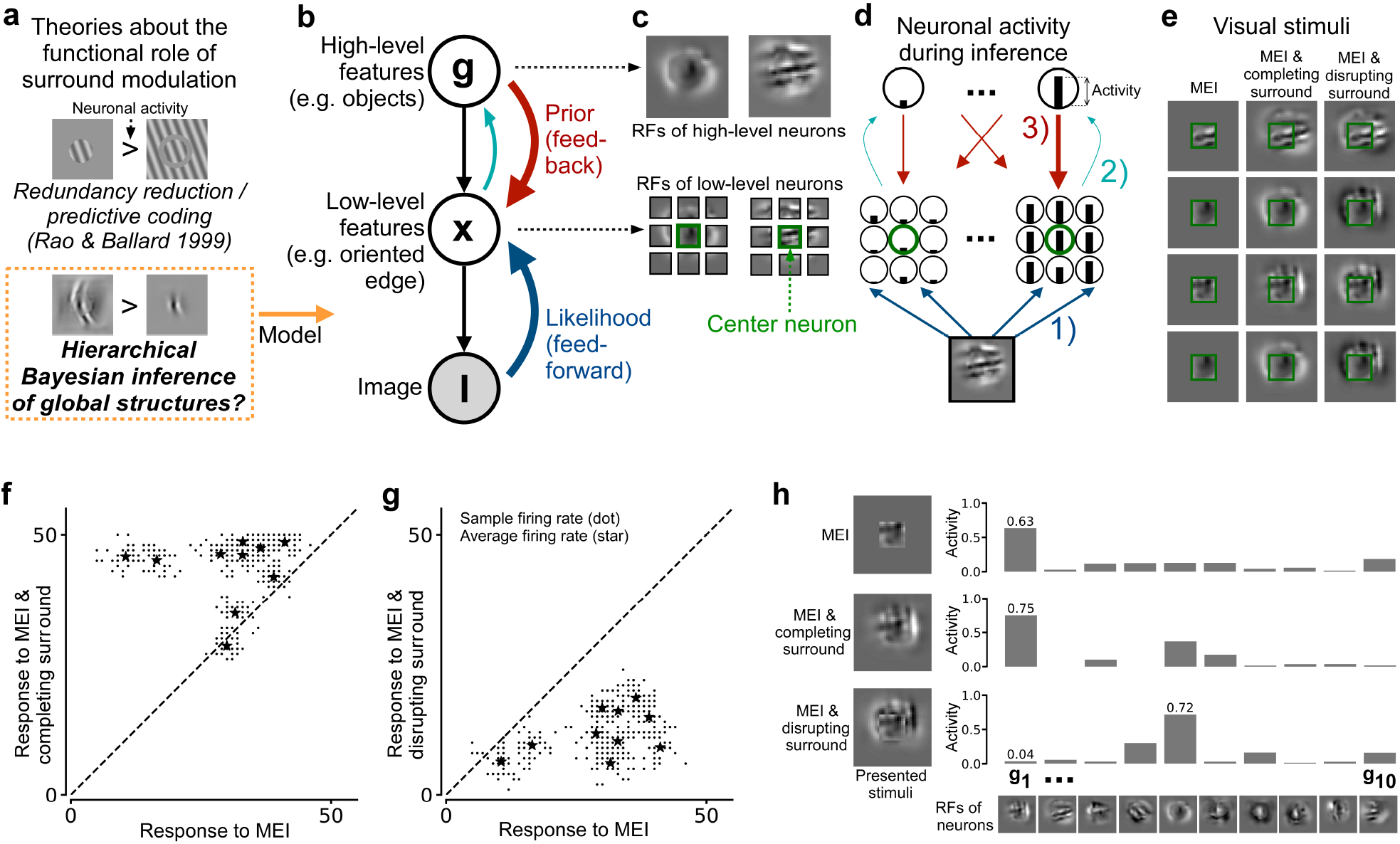
Explaining observed center-surround effects by Bayesian inference. **a**, Schematic illustrating theories about the functional role of surround modulation. Weaker suppression by ortho-than iso-oriented gratings in the surround has been linked to redundancy reduction and efficient coding. Here, we propose that pattern completion and disruption by the excitatory and inhibitory surround, respectively, could emerge from perception through hierarchical Bayesian inference of global features. **b**, Schematic illustrating the visual system as a generative model of the stimulus, **I. g** represents high-level features (e.g. objects), and **x** represents low-level features (e.g. oriented edges). All three variables are multidimensional. Shaded circles denotes observed, open circles inferred variables. Inferring the posterior over **x** entails combining likelihood (feedforward), and prior expectations driven the belief about which high-level features are present (feedback). **c**, RFs of both **g** and **x** neurons. RFs of **g** neurons represent full objects, spanning all pixels in the image, **I**. RFs of **x** neurons represent local features. Some **x** represent the center (green border) and others the surround. **d**, Illustration of information flow during inference. Each dimension in **g** represents a neuron in visual areas downstream to V1, each encoding the presence of an object. Each dimension in **x** represents a neuron in V1, each encoding the presence of a specific local feature. Feedback signals from a single *gi* boost compatible *xj*. **e**, Four example MEIs shown with their corresponding completing surround, and disrupting surround (see Methods section for details on how they are constructed). **f-g**, Scatterplots of simulated neural activity during inference of MEI vs completing and disruptive surrounds respectively. Our model consisted of 10 textures/objects (**g**) and 90 V1 neurons, (**x**), with the RFs of 10 neurons at the center of the image, and 80 neurons with RFs in their surround. Scatterplots show responses from all 10 center neurons. Each dot (*) corresponds to the firing rate during a single simulated trial and each star () corresponds to one center neuron’s average firing rate. **h**, Marginal response distribution over higher-ordered **g** neurons for an example experiment.

Our hierarchical generative model is similar to ones previously proposed (Haefner et al., 2016; Bányai et al., 2019) (Fig 7b,c). In the generative model, we assume that V1 neurons represent the presence of local spatial features (*x*) neurons and higher order areas represent the presence of objects or larger textures (*g*). The goal of this model visual system is to infer the presence of local spatial features and hierarchically, the presence of objects or textures. In other words, the goal is to perform joint probabilistic posterior inference over **x** and **g** given an image. As a consequence, inference *p*(**x, g**|**I**)*∝p*(**x**|**g**) *p*(**I**|**x**) over the intermediate variables **x** — representing V1 neurons — and global variables **g** — representing higher order neurons — combines two types of information: feedforward *p*(**I**|**x**) from the input image **I**, and feedback *p*(**x**|**g**) from higher level areas reflecting expectations resulting from the current belief about which global feature is present (Fig 7b-d).

To quantify the center-surround interactions in this model, we presented the following three sets of stimuli tailored to the V1 neurons whose RFs are located in the center of visual space (Fig. 7c, RFs with green border): (1) the MEI of the V1 neurons, (2) the MEI with a spatially completing pattern in the surround, and (3) the MEI with a spatially disrupting pattern in the surround (Fig. 7e). These three conditions match the pattern completion and disruption that characterize the contextual modulations we found in mouse and primate visual cortex. For each stimulus condition, we performed joint posterior inference in the generative model, i.e., computed the posterior distribution *p* (**g, x**|**I**) and obtained the responses of both **g** and **x** neurons. Subsequently, we compared the responses of the center-aligned V1 neurons to their respective MEIs with the responses elicited by (1) the MEIs with the completing surround and (2) the MEIs with the disrupting surround. The model responses reproduced our key experimental results (Fig. 7f-g): the MEI with the spatially completing surround drives the center-aligned V1 neurons stronger than its MEI alone, and the MEI with the spatially disrupting stimulus inhibits the responses of the neurons compared to the MEI presented alone.

The key driver of excitation and inhibition in our probabilistic model is the top-down signal resulting from beliefs about the presence or absence of large-scale features. A V1 neuron’s response is boosted when its feedforward input is congruent with the brain’s beliefs about what that input should be. This belief is strongest when image center and surround are congruent (completing surround) and indicative of the same global feature. On the other hand, it is weakened when the surround is incongruent with the center (disrupting surround). In particular, when only the MEI is present, the corresponding global feature may be inferred to be present with an intermediate probability (0.63 in the example in Fig 7h, top row). When a congruent surround is added, this probability increases (0.75, Fig 7h, middle row). However, when an incongruent surround is added, this probability decreases (0.04, Fig 7h, bottom row). Consequently, the activity of the corresponding V1 neuron is enhanced for the congruent surround, and suppressed for the incongruent surround, relative to the MEI-only condition.

## Discussion

Our study discovered a novel rule of surround modulation in primary visual cortex: Completion (or extension) of naturalistic visual spatial patterns in the center RF governed surround excitation, whereas disruption (or termination) of center features produced inhibition. The non-linearity of neuronal responses to natural images, which reside in a high-dimensional space, has made it challenging to accurately characterize the center RF properties and to model the interactions with the RF surround in the context of natural visual inputs. Our accurate digital twin models allowed us to capture the non-linearity both within and beyond the center RF, and to predict the best modulating stimuli in the surround, without parametric assumptions about their underlying statistical structure. We verified the predictions from the model experimentally in a closed-loop manner. Our results demonstrate that contextual modulation in mouse primary visual cortex is driven by pattern completion and disruption shaped by natural image statistics. Additionally, our results suggest that a similar mechanism of excitatory surround pattern completion is also present in the macaque primary visual cortex. This type of surround facilitation by congruent structures emerged within a simple hierarchical model that modulates neuronal responses based on prior knowledge of the world, i.e. natural scene statistics. This may potentially enhance the encoding of prominent features in the visual scene, such as contours and edges, especially when the sensory input is noisy and uncertain.

### Relationship between surround modulation and stimulus statistics

Previous studies using oriented stimuli such as gratings and bars have explored spatial patterns of contextual modulation in the primary visual cortex of monkeys (Allman et al., 1985; Levitt and Lund, 1997; Kapadia et al., 1999; Sceniak et al., 1999; Cavanaugh et al., 2002b,c; Nassi et al., 2013; Nurminen et al., 2018; Michel et al., 2018; Knierim and Van Essen, 1992; Polat et al., 1998). These investigations predominantly identified suppression, particularly from congruent surround stimuli, as the dominant modulation form of surround modulation, with the strength of suppression waning as surround stimulus congruency decreases (Knierim and Van Essen, 1992; Kapadia et al., 1999). Our results in the macaque V1 model are consistent with these findings, confirming that suppression is the predominant effect of iso- and ortho-oriented gratings in the surround RF. However, it is important to note that the surround modulation dynamics vary significantly with the stimulus configuration. Previous studies have shown that at lower contrast and with specific arrangements such as co-linear bars adjacent to the center RF instead of full-field gratings, congruent stimuli in the RF surround can elicit excitation (Polat et al., 1998; Lee and Nguyen, 2001). As a whole, the literature on surround modulation in primate visual cortex suggests that details of the stimulus like contrast, size and location greatly influence both the strength as well as the effect of surround modulation on neuronal responses (reviewed in Angelucci et al., 2017).

So far, the spatial patterns driving surround excitation versus inhibition in mouse V1 are less conclusive compared to primates. Some previous studies have reported suppression and facilitation of mouse V1 neurons by congruent and incongruent parametric surround stimuli (Keller et al., 2020a; Self et al., 2014), respectively, consistent with the results in primates. However, there seems to be a large variability across neurons, where surround stimuli that have the same orientation as the center stimulus can be either excitatory or inhibitory (Samonds et al., 2017) and different orientations of the surround relative to the center can be excitatory (Keller et al., 2020b). Here, we have confirmed those results by using iso- and ortho-oriented drifting grating stimuli. In part, this variability across neurons might be related to the fact that parametric stimuli like gratings and bars drive mouse V1 neurons sub-optimally, due to the fact that mouse V1 neurons are selective for more complex visual features (Walker et al., 2019). It is well established that contextual modulation depends on the center stimulus features (Knierim and Van Essen, 1992; Kapadia et al., 1999) and it might therefore be critical to condition surround stimuli on the optimal stimulus in the center RF, corresponding to the MEI (Walker et al., 2019).

Our results, obtained using naturalistic stimuli and a data-driven approach that minimizes strong assumptions about stimulus selectivity, revealed a novel principle of surround modulation in the mouse primary visual cortex. We discovered that the most excitatory surround stimuli are congruent with the optimal center stimuli, thus completing patterns according to natural image statistics, as shown using a generative diffusion model. Conversely, the most inhibiting surround stimuli are incongruent, thereby disrupting these patterns. Our findings establish a consistent rule of pattern completion and disruption, which leads to surround facilitation and suppression. We further demonstrated that this principle also applies to macaque V1, particularly in the near surround where excitatory surround images more frequently complete patterns compared to inhibitory ones, reminiscent of collinear facilitation reported using bars and gratings (Levitt and Lund, 1997; Polat et al., 1998; Keller et al., 2020b). However, un-like prior research which indicated facilitation for very specific stimulus configurations, our study proposes a new universal rule—pattern completion—that consistently leads to surround facilitation, both in mouse and primate V1 neurons, and is related to the spatial statistics of natural images that go beyond collinearity. For example, in mouse V1, the most exciting images are not typically Gabors, thus making it unclear how collinearity would apply or what an optimal surround would be when the optimal center stimulus is a texture, for example, or a corner. The use of an image synthesis approach revealed that the non-parametric excitatory surround patterns of macaque V1 neurons incorporate complex patterns with varying orientations, often reflective of natural scene configurations. This suggests that the facilitation by collinear structures might represent a simplification of our newly identified rule, underscoring the effectiveness of the digital twin model in conducting exhaustive in-silico experiments. These experiments explore both non-parametric and parametric stimuli, helping to reconcile the diverse effects of contextual modulation observed under different stimulus conditions.

Overall, our findings complement previous studies that noted suppressive effects from iso- and ortho-oriented surround gratings in both mouse and macaque V1, which we replicated and analyzed in our experiments. Our work enhances the understanding that contextual modulation is critically influenced by the statistical properties of stimuli. It shows that for a non-parametric approach, surround modulation is driven by pattern completion and disruption. This mechanism, shaped by natural scene statistics in mice, is also suggested to be present in macaques.

A recent study by Pan et al. (2023) utilized a similar non-parametric approach to synthesize surround images for hidden units in artificial neural networks, finding that congruent spatial patterns in the center and surround are most suppressive. This appears to contradict our results, suggesting a potentially intriguing divergence between natural visual systems and current artificial neural networks. Notably, the findings reported by Pan et al. (2023) vary significantly depending on the network layer and its architecture. Exploring these differences between various artificial neural network architectures and digital twins of the brain represents a promising direction for future research, and promises to uncover universal principles of visual information processing conserved across both animal species and artificial vision systems.

### Circuit-level mechanism of contextual modulation in visual cortex

Mechanistically, surround suppression in V1 can be partially accounted for by feedback projections from higher visual areas. In monkeys, inactivation of feedback from V2 and V3 reduces surround suppression induced by large grating stimuli (Nassi et al., 2013; Nurminen et al., 2018) and also results in an increase in RF size (Nurminen et al., 2018). In mice, feedback from higher visual areas also strongly modulates V1 responses to stimuli in the RF center and even elicits strong responses without any stimulation of the center, thereby creating a feedback RF (Keller et al., 2020b; Shen et al., 2022). The cellular substrate of surround modulation has been predominantly studied in mice, benefiting from genetic tools for cell-type specific circuit manipulations. Different types of inhibitory neurons have been identified as key players of surround modulation, including somatostatin (SOM)- and vasoactive intestinal peptide (VIP)-expressing cells, which inhibit each other as well as excitatory V1 neurons and are further modulated by feedback (Adesnik et al., 2012; Keller et al., 2020a; Shen et al., 2022). Based on these results, surround suppression in mouse V1, and likely primate V1, is dependent on the exact balance between the excitatory input from feedforward and feedback projections and the inhibitory inputs from locally present inhibitory neuron types.

To further elucidate surround modulation of individual visual neurons in relation to local and long-range network connectivity, recent advancements in functional connectomics offer significant opportunities. These advances combine large-scale neuronal recordings with detailed anatomical information at the scale of single synapses. Utilizing the MICrONS functional connectomics dataset (MICrONS Consortium et al., 2021) and its functional digital twin (Wang et al., 2023), we investigated circuit-level mechanisms that could underlie the pattern-completion governed surround facilitation observed in our data. This dataset encompasses responses from over 75,000 neurons to natural movies, along with the reconstructed sub-cellular connectivity of these cells from electron microscopy data. Our analysis identified ‘like-to-like’ anatomical connections among neurons with similar feature selectivity but minimal RF overlap, which likely facilitates the completion of naturalistic patterns observed in excitatory surround images. Furthermore, our modeling results suggest that inhibitory surround modulation may be driven by the collective activity of a functionally diverse group of neurons, aligning with earlier studies (DeAngelis et al., 1992; Morrone et al., 1982) and pointing to an additional circuit motif underlying surround suppression in the visual cortex. As connectomics proofreading efforts for the MICrONS dataset proceed, aiming to reconstruct the connectome among tens of thousands of excitatory and inhibitory neurons across various cortical layers and visual areas, we anticipate gaining a much more comprehensive understanding of the circuit-level mechanisms behind contextual modulation. This progression will enable us to extend our connectivity analysis from excitatory neurons within V1 to higher cortical areas to explore feedback projections, and to interneurons to examine feature-specific inhibitory inputs to projection neurons, akin to studies performed on the fly visual system connectome (Sebastian Seung, 2024). The creation of a functional digital twin of the MICrONS dataset (Wang et al., 2023) and our demonstration of its utility in studying the circuit-level mechanisms of neuronal computations, showcased here for center-surround interactions, promise significant progress in understanding both structure and function of neuronal circuits.

### Theoretical implications of surround facilitation

Here, we demonstrated that surround facilitation is a prominent feature of contextual modulation in the primary visual cortex, thereby highlighting that center-surround interactions cannot simply be explained by suppression of sensory responses. Importantly, excitatory surround images with the optimal center stimulus exhibited a high representational similarity with natural images, indicating that congruent patterns frequently present in natural scenes (Geisler et al., 2001; Sigman et al., 2001) strongly drive neuronal responses, through excitatory surround pattern completion. Excitation by congruent surround structures relative to the center may be explained by preferential long-range connections between neurons with co-linearly aligned RFs described in mice (Iacaruso et al., 2017) and higher mammals (Bosking et al., 1997; Schmidt et al., 1997; Sincich and Blasdel, 2001) and might serve perceptual phenomena like edge detection, contour integration and object grouping observed in humans and primates (Kapadia et al., 1995; Geisler et al., 2001).

Our empirical results of surround facilitation are surprising in light of a long line of theoretical work that explains sensory responses using principles like redundancy reduction (Barlow et al., 1967) or predictive coding (Rao and Ballard, 1999). The idea that neurons should minimize redundancy has given rise to contrast normalization models (Schwartz and Simoncelli, 2001) that were recently expanded to a flexibly-gated center-surround normalization model (Coen-Cagli et al., 2015) most relevant to our data. The key idea behind the latter model is to only normalize (typically reduce) center activation when the surround is similar, and otherwise ignore the surround. This proposal cannot explain our empirical findings. Analogously, predictive coding proposes that neuronal activity reflects prediction errors, and that therefore the center activation should be lower when it can be well predicted from the surround (Rao and Ballard, 1999; Keller and Mrsic-Flogel, 2018) – again in contradiction to our finding that excitatory surrounds appear to “complete” the center stimulus, and frequently occur in natural scenes.

In contrast, our results are consistent with an alternative framework for understanding sensory neurons: perceptual (Bayesian) inference (Von Helmholtz, 1867; Knill and Richards, 1996). In this model, sensory responses calculate beliefs about latent variables within a hierarchical structure, where higher-level variables represent broader, more complex image features and act as priors for lower-level variables. These lower variables represent specific parts of the image and receive feedback from higher levels (Lee and Mumford, 2003). In such a model, global image structure can increase or decrease responses of neurons with localized RFs, depending on whether the global structure increases or decreases the probability of the local feature being present in the image (Haefner et al., 2016; Bányai et al., 2019; Lange and Haefner, 2022). In fact, our probabilistic model which qualitatively reproduces our empirical findings is an example of such a model. Our approach of characterizing contextual modulation in a data-driven way for arbitrary stimuli, without any assumptions about neuronal selectivity, has revealed a novel relationship between surround modulation and natural image statistics that challenges classic theories of redundancy reduction and predictive coding, instead providing evidence a contextual modulation expected from models of hierarchical inference in which neurons represent beliefs about the outside world.

It is possible that different computational objectives may co-exist and operate under different input regimes. At high certainty (e.g., high contrast), the efficiency achieved by redundancy reduction might be most important. Conversely, in high uncertainty scenarios (e.g., low contrast), maximizing information by incorporating prior knowledge of the world by Bayesian inference might be more advantageous.

## Materials and Methods

### Animals and surgical preparation

All experimental procedures complied with guidelines of the NIH and were approved by the Baylor College of Medicine Institutional Animal Care and Use Committee (permit number: AN-4703), expressing GCaMP6s in cortical excitatory neurons. Mice used in this study (n=14, 7 males and 7 female, aged 2.5 to 6 month) were heterozygous crosses between Ai162 and Slc7a7-Cre transgenic lines (JAX #031562 and #023527, respectively). To expose V1 for optical imaging, we performed a craniotomy and installed a window that was 4mm in diameter and centered at 3mm lateral to midline and 2mm anterior to lambda (Reimer et al., 2014; Froudarakis et al., 2014). Mice were housed in a facility with reverse light/dark cycle to ensure optimal alertness during the day when experiments were performed.

### Neurophysiological experiments and data processing

We recorded calcium signals using 2-photon imaging with a mesoscope (Sofroniew et al., 2016) which was equipped with a custom objective (0.6 numerical aperture, 21 mm focal length). The imaging fields of each recording were 630×630 µm^2^ per frame at 0.4 pixels µm^-1^ xy resolution and were positioned in the center of V1 according to the retinotopic map (Fig. 1b). Z resolution was 5 µm with a total of ten planes from *−*200*µm* to *−*245*µm* relative to cortical surface. The laser power increased exponentially as imaging plane moved farther from the surface according to:

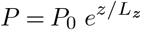

Here *P* is the laser power used at target depth *z, P*_0_ is the power used at the surface (19.71 mW ±4.68, mean ±standard deviation), and *L*_*z*_ is the depth constant (220 µm). The highest laser output was of 54.79 mW ±13.67 and was used at approximately 240 µm from the surface. Most scans did not require more than 50 mW at maximal depth, except for one mouse where the average laser power at the deepest scanning field was 82.03 mW.

For each animal, we first performed retinotopic mapping across the whole cranial window to identify the border of V1 (Fig. 1b and c; Schuett et al., 2002). At the beginning of each imaging session, we measured the aggregated population RF to ensure precise placement of the monitor with regard to the imaging site. We used stimuli consisting of dark (pixel value=0) square dots of size 6 degrees in visual angle on a white background (pixel value=255). The dots were randomly displayed at locations on a 10 by 10 grid covering the central region of the monitor and at each location the dot was shown for 200 ms and repeated 10 times over the whole duration of dot mapping. The mean calcium signal was deconvolved and averaged across repeated trials to produce the population RF. The monitor was placed such that the population RF was centered on the monitor.

The full two-photon imaging processing pipeline is available at (https://github.com/cajal/pipeline). Briefly, raster correction for bidirectional scanning phase row misalignment was performed by iterative greedy search at increasing resolution for the raster phase resulting in the maximum cross-correlation between odd and even rows. Motion correction for global tissue movement was performed by shifting each frame in x and y to maximize the correlation between the cross-power spectra of a single scan frame and a template image, generated from the Gaussian-smoothed average of the Anscombe transform from the middle 2000 frames of the scan. Neurons were automatically segmented using constrained non-negative matrix factorization, then traces were deconvolved to extract estimates of spiking activity, within the CalmAn pipeline (Giovannucci et al., 2019). Cells were further selected by a classifier trained to separate somata versus artifacts based on segmented cell masks, resulting in exclusion of 8.1% of the masks.

A 3D stack of the volume imaged was collected at the end of each day to allow registration of the imaging plane and identification of unique neurons. The stack was composed of two volumes of 150 planes spanning from 50 µm above the most superficial scanning field to 50 µm below the deepest scanning field. Each plane was 500×800 µm, together tiling a 800×800 µm field of view (300 µm total overlap), and repeated 100 times per plane.

### Visual stimulation

Visual stimuli were displayed on a 31.8 ×56.5 cm (height × width) HD widescreen LCD monitor with a refresh rate of 60 Hz at a resolution of 1080 × 1920 pixels. When the monitor was centered on and perpendicular to the surface of the eye at the closest point, this corresponded to a visual angle of 2.2^*◦*^/cm on the monitor. We recorded the voltage of a photodiode (TAOS TSL253) taped to the top left corner of the monitor to measure the gamma curve and luminance of the monitor before each experimental session. The voltage of the photodiode is linearly correlated with the luminance of the monitor. To convert from photodiode voltage to monitor luminance, we used a luminance meter (LS-100 Konica Minolta) to measure monitor luminance for 16 equidistant pixel values from 0-255 while recording the photodiode voltage. The gamma value for experiments in this paper ranged from 1.751 to 1.768 (mean = 1.759, standard deviation = 0.005). The minimum luminance ranged from 0.23 cd/m^2^ to 0.97 cd/m^2^ (0.49 ± 0.25, mean ± standard deviation), and the maximum ranged from 84.11 cd/m^2^ to 86.04 cd/m^2^ (85.07 ± 0.72, mean ± standard deviation).

### ImageNet stimulus

Natural images were randomly selected from the ImageNet database (Deng et al., 2009), converted to gray scale, and cropped to the monitor aspect ratio of 16:9. To probe center-surround interactions, we modified the images using a circular mask that was approx. 48 degrees in visual angle in diameter with smoothed edges. The mask radius was defined as fraction of monitor width, i.e. *r*_aperture_ = 1 means a full-field mask. We used *r*_aperture_ = 0.2

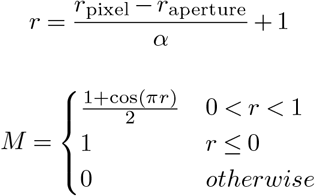

where *M* is the mask, *r* is the radius, and *α* is the width of the transition. We presented 5,000 unique natural images without repetition during each scan, half of which were masked. We also presented the same 100 images repeated 10 times as full-field and 10 times as masked. The 100 images that were repeated were conserved across experiments, while the unique images varied across scans. Each trial consisted of one image presented for 500 ms with a preceding blanking period of 300 - 500 ms (randomly determined per trial).

### Eye tracking

A movie of the animal’s eye and face was captured throughout the experiment. A hot mirror (Thorlabs FM02) positioned between the animal’s left eye and the stimulus monitor was used to reflect an IR image onto a camera (Genie Nano C1920M, Teledyne Dalsa) without obscuring the visual stimulus. The position of the mirror relative to the camera was manually adjusted if necessary per session to ensure that the camera focuses on the pupil. The field of view was manually cropped for each session. The field of view contained the left eye in its entirety, 282-300 pixels height × 378-444 pixels width at 20 Hz. Frame times were time stamped in the behavioral clock for alignment to the stimulus and scan frame times.

Light diffusing from the laser during scanning through the pupil was used to capture pupil diameter and eye movements. A DeepLabCut model (Mathis et al., 2018) was trained on 17 manually labeled samples from 11 animals to label each frame of the compressed eye video with 8 eyelid points and 8 pupil points at cardinal and intercardinal positions. Pupil points with likelihood >0.9 (all 8 in 93%± 8% of frames) were fit with the smallest enclosing circle, and the radius and center of this circle was extracted. Frames with <3 pupil points with likelihood >0.9 (0.7%± 3% frames per scan), or producing a circle fit with outlier >5.5 standard deviations from the mean in any of the three parameters (center x, center y, radius, <1.3% frames per scan) were discarded (total <3% frames per scan). Trials affected by gaps in the frames were discarded (<2% trials for all animals except one, where the animal’s eye appeared irritated).

### Registrations of neurons in 3D stack

We densely sampled the imaging volume to avoid losing cells due to tissue deformation from day to day. Therefore, some cells were recorded in more than one plane. To select unique cells, we subsampled our recorded cells based on proximity in 3D space. Each functional scan plane was independently registered to the same 3D structural stack. Specifically, we used an affine transformation matrix with 9 parameters estimated via gradient ascent on the correlation between the sharpened average scanning plane and the extracted plane from the sharpened stack. Using the 3D centroids of all segmented cells, we iteratively grouped the closest two cells from different scans until all pairs of cells are at least 10 µm apart or a further join produces an unrealistically tall mask (20 µm in z). Sequential registration of sections of each functional scan into the structural stack was performed to assess the level of drift in the z dimension. The drift over the 2 to 2.5 hour recording was 4.70± 2.64, and for most of them the drift was limited to <5 µm.

### Model architecture and training

The convolutional neural network used in this study consisted of two parts: a core and a readout. The core captured the nonlinear image representations and was shared among all neurons. The readout mapped the features of the core into neuronal responses and contained all neuron specific parameters.

### Core

To get a rich set of nonlinear features, we used a deep CNN as our core. We used a CNN with 3 layers and 32 feature channels per layer as previously described in (Walker et al., 2019). These architectures were chosen with a hyperparameter search, with the objective of maximizing a validation score (see **Training and evaluation**). Each of the 2D convolutional layers was followed by a batch normalization layer and an ELU non-linearity.

### Readouts

The goal of the readout was to find a linear-nonlinear mapping from the output of the last core layer Φ(**x**) to a single scalar firing rate for every neuron. We used a pyramid readout, as described in Sinz et al. (2018). We computed a linear combination of the feature activations at a spatial position, parameterized as (*x, y*) relative coordinates (the middle of the feature map being (0, 0)). We then passed these features through a linear regression and a non-linearity to obtain the final neuronal responses.

### Training and evaluation

Natural images in the training, validation and test sets were all Z-scored using the mean and standard deviation of the training set. The mean and standard deviation for the cropped natural images were weighted by the mask used to crop the images to avoid artificially lowering the mean and standard deviation due to large gray areas in the cropped images.

The networks were trained to minimize Poisson loss 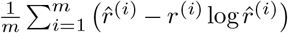 where *m* denotes the number of neurons, 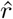 the predicted neuronal response and *r* the observed response. We implemented early stopping on the correlation between predicted and measured neuronal responses on the validation set: if the correlation failed to increase during 10 consecutive epochs through the entire training set, we stopped the training and restored the best performing model over the course of training. After each stopping, we either decreased the learning rate or stopped training altogether if the number of learning-rate decay steps was reached. Network parameters were optimized via stochastic gradient descent using the Adam optimizer. Once training completed, the trained network was evaluated on the validation set to yield the score used for hyper-parameter selection.

### MEI and surround image generation

Because our neuronal recordings were performed with dense sampling (Z spacing = 5*µm*), we first needed to select unique neurons. We registered the planes of the functional experiments to the stack of the volume (see **Registration of neurons in 3D stack**) and identified unique neurons.

Then, we optimized the MEIs and the surround images in two steps.

### MEI generation

We used regularized gradient ascent by solving the optimization problem defined as

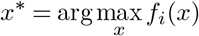

on our trained deep neural network models to obtain a maximally exciting input image for each neuron, given by *x*

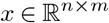

(Walker et al., 2019). We initialized with a Gaussian white noise image. In each iteration of gradient ascent, we showed the image to the model and calculated the gradients of the image w.r.t. the model activation of a single neuron. We then blurred the obtained gradient with Gaussian blurring, with a Gaussian sigma of 1 pixel. Following this, we updated the image with the resulting gradients. Finally, we calculated the standard deviation of the resulting image and rescaled its contrast to match a fixed RMS contrast constraint of 0.05 (in z-scored response space). The contrast constraint was chosen to minimize the number of pixel values falling outside the range 0 and 255, which are the lower and upper bound for pixel values displayed on the monitor. The RMS contrast constraint of 0.05 for the full-field MEI images resulted in a RMS contrast of 12.15 ±1.35 in 8-bit input space (0 to 255 pixel values) within the MEI mask. For a subset of experiment, we used a RMS contrast constraint of 0.1, resulting in a RMS contrast of 22.23 ±3.38 in 8-bit space within the MEI mask. We used the Stochastic Gradient Descent (SGD) optimizer with step size=0.1 and ran each optimization for 1,000 iterations.

### Surround image generation

A tight mask (ranging between 0 and 1) around the MEI was computed by thresholding (see below) which we used to define the ‘center’ and set it apart from the ‘surround’ during the next step of optimization. By applying the inverse MEI mask to the target image x, we optimized the surrounding area in the image by allowing more contrast (RMS contrast = 0.1) outside of the MEI mask.

To define the center stimuli, we computed a mask around the MEI for each neuron by thresholding at 1.5 standard deviations above the mean. We then blurred the mask with Gaussian *σ* = 1 pixel. We initialized an image with Gaussian noise and cropped out the center of this image using the MEI mask and added the MEI at a fixed contrast = 0 .05. We s et the contrast for the area outside of the mask to 0.1. For the high contrast experiments, the surround contrast was set to 0.2. A gradient was computed on the modified image and we blurred the gradient with a Gaussian *σ* = 1. We used the same SGD optimizer to update the image at each iteration.

Only pixels outside of the MEI mask were updated during optimization (illustrated in Fig. 2a). We set the full-field image contrast to an arbitrary value within the training image regime (0.1) to prevent the pixel values from getting out of range and this step was not differentiable. At the end of each iteration, we normalized the contrast in the center and the surround again to reach the optimal stimulus with correct contrast (MEI=0.05, surround=0.1 or MEI=0.1, surround=0.2). We repeated these steps for 1,000 iterations. To generated the extended mask for the MEI used in Suppl. Fig. 4, we set the value between 1 and 0.001, i.e. in the blurred area, in the original mask to 1 and blurred the new mask with the same Gaussian filter that was applied to the MEI mask. We applied the extended mask to the surround images to produced a new set of masked surround images that were slightly smaller than the original ones, and tested surround modulation restricted only to the “near” surround region.

### Closed-loop experiments

#### Selection of neurons for closed-loop

We ranked the neurons recorded in one experiment based on response reliability and model performance (test correlation). Specifically, we correlated the leave-one-out mean response with the remaining single-trial response across repeated images in the test set to obtain a measurement of neuronal response reliability. We then computed an averaged rank score of each neuron from its reliability rank and model test correlation rank. After removing duplicate neurons following the procedure described above, we selected the top 150 neurons according to the averaged rank of the correlation between predicted response and observed response averaged over repeats and the correlation between the leave-one-out mean response of repeated test trials to the left-out test trial response for closed-loop experiments. Please note that due to this selection process, our conclusions are limited to the neurons in the dataset that demonstrated reliable responses and were accurately predicted by our model.

#### Stimulus presentation

We converted the images generated by the model back to pixel space by reversing the Z-score step with the stats of the training set. Each image was repeated 40 times. We shuffled all the images with repeats across different classes (MEI, excitatory, inhibitory and outpainted surrounds and contrast-matched MEI, masked surround controls) and presented them at random orders. Each trial consisted of one image presented for 500 ms with a preceding blanking period of 300 - 500 ms (randomly determined per trial).

#### Matching neurons across experiments

We matched neurons from different experiments according to the spatial proximity in the volume of the same anatomical 3D stack. Each functional scan plane was registered to the 3D stacks collected after each day’s experiment. We chose the neurons that had the highest matching frequency across all stacks, and included them as a valid neuron in the closed-loop analysis.

#### Estimation of center RF size

To measure to size of the minimum response field (MRF) for each neuron, we presented stimuli consisting of circular bright (pixel value=255) and dark (pixel value=0) dots of size 7 degrees in visual angle on a gray background (pixel value=128) in conjunction with natural image stimuli. The dots were randomly shown at locations on a 9 by 9 grid covering 40% of the monitor in the center along the horizontal edge, and at each location, the dot was shown for 250 ms and repeated 16 times. The responses were averaged across repeats, and a 2D Gaussian was fitted to the On and Off response maps, respectively. The size of the MRF was measured as the largest distance between points on the border of the 2D Gaussian at 1.5 standard deviations away for both On and Off responses.

To estimate the size of the MEIs and the excitatory and inhibitory surround, we first computed the mask for each image as described in section **MEI and surround image generation**. The size was computed in pixels as the longest distance between points on the border of the mask. The size was converted to degrees in visual angle according to the ratio between pixel and degrees in visual angle.

#### Exciting natural image patches and natural surrounds

All natural images in the ImageNet dataset were first Z-scored with the mean and standard deviation of the training dataset. We then cropped the images with the MEI masks and normalized to match the contrast of the MEI within the mask. The images were presented to the model to get the predicted response. Images that elicited activations above 80% of MEI activation were chosen as the maximally exciting natural image patches. Images used to train the specific model were removed from this collection. For neurons with more than 10 maximally exciting natural image patches, we replaced the center of the natural image with the MEI and included the surround region of the natural image to the same extend as the average size of the excitatory and the inhibitory surround.

#### Representational similarity

The maximally exciting natural image patches of a neuron plus the surround of the same image were normalized to the same contrast as the excitatory and the inhibitory surround images and were presented to the model. The excitatory and the inhibitory surround images were cropped with the average mask of the two to match the size, contrast-adjusted and presented to the model. The activation of all neurons in the model were taken as an approximation of the given image in “representational space”. We computed Pearson correlation between a natural image patch with surround and an image of the MEI with either excitatory or inhibitory surround. The Pearson correlation is an estimation of ‘representational similarity’.

#### Diffusion outpainted surround images

We performed outpainting by drawing samples from the posterior *p*(*y*|*x*^*∗*^), where *x*^*∗*^ is the MEI and *y* is the outpainted image. To generate samples from this posterior we use energy guided diffusion (Pierzchlewicz et al., 2023), where the score of the posterior is defined as:

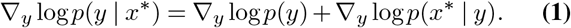

The prior is defined by the ablated diffusion model Dhariwal and Nichol (2021) *ε*_*θ*_(*y*) pre-trained on ImageNet acting as a natural-image prior. The likelihood is defined by the energy

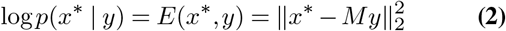

where *M* is the MEI mask. The images generated by the diffusion model are square, thus we first increased the resolution of the MEI image from 36×64 to 144×256 by bi-linear interpolation and then squarified by padding it with zeros to achieve 256×256. The final sample is then cropped to 144×256 and down-scaled to 36×64 and masked by the excitatory or inhibitory surround mask.

### In-silico analysis of macaque V1 neurons

#### Macaque V1 digital twin model

We used a previously published dataset (see details in (Safarani et al., 2021; Cadena et al., 2023; Baroni et al., 2023)) for model training. In brief, we measured the spiking activity of individual V1 neurons in two awake, fixating rhesus macaques using a 32-channel linear array spanning multiple cortical layers, in response to tens of thousands of grayscale natural images, covering 6.7^*◦*^ visual angle, presented in sequence over many trials. These images were sampled uniformly from the ImageNet (**?**) dataset and displayed for 120 ms each without interleaving blanks. Most of these images were shown only once (train-set) while a selection of 75 images was repeated multiple times (test-set). We isolated 458 V1 neurons from 32 sessions at eccentricities 2–3°. We centered the stimuli on the population receptive field of the neurons. Finally, we obtained image–response pairs by extracting spike counts in the window 40–160 ms after image onset. With these image–response pairs, we fitted our models. Before presenting the images to the model, we effectively cropped the images down to the central 2.67°, corresponding to 93 by 93 px.

Like the digital twin for all the mouse models described in this study, the neural predictive model for the macaque V1 data consisted of two main parts: A pre-trained core that computes image embeddings, i.e. a shared feature map given an input images, and a readout that maps these features to the neuronal responses of a single neuron. As a core, we selected ConvNext-v2-tiny (Woo et al., 2023), a recently published convolutional neural network model trained on ImageNet. We used the original neural network weights obtained from the transformers library of huggingface (Wolf et al., 2019) and performed a hyperparameter search, which output layer resulted in the best predictive performance, which was *stages-1-layers-0*. As readout, we fit a Gaussian read-out, described in detail in (Lurz et al., 2021), to transform the core feature map into a scalar neural response for each recording channel. Finally, a neuron-specific affine projection with ELU non-linearity gives rise to the scalar predicted neuronal activity. The model is being trained by minimizing the Poisson loss between recorded and predicted neuronal activity, identically to the procedures described in Willeke et al. (2023). Here, we first freeze the core weights and train the readout for 20 epochs. Then, we reduce the initial learning rate from 0.001 to 0.0001 and optimize the weights of both the convnext core and readout, using the AdamW optimizer (Loshchilov and Hutter, 2017) for a total of 200 epochs. We trained n=5 models with different random seeds and used these as an ensemble by averaging the predictions of each model. For all subsequent analyses, we used the ensemble model and refer to it simply as model. The model performance, measured as the correlation between model predictions and the average neuronal response across repeats, was 0.74, evaluated on the held-out test set of 75 test images, out-performing the best ResNet-based models (He et al., 2016) which achieved a correlation of 0.66 (Cadena et al., 2023) and the best purely data-driven, i.e. end-to-end trained model (Baroni et al., 2023) with a correlation of 0.72.

#### Classical grating experiments

We conducted a list of insilico experiments on macaque V1 neurons. To identify RF position and size we performed a sparse noise experiment. Stimuli consisted of white or black squares of 4×4 pixels (corresponding to 0.11×0.11 degrees) on a mid-scale grey background. In order to obtain the RF for each neuron, we first computed a polarity agnostic version of the stimuli (mapping black squares to white squares). Then we computed a weighted average of the polarity-agnostic stimuli according to responses after subtraction of the baseline response (response to a midscale-grey background only). In this way, we obtained an RF estimate showing areas of excitation and suppression. Then, we clipped the output pixel values below 0, in order to remove the suppression effect. Lastly, we fitted the output with a 2D Gaussian. We estimated the neuron’s RF position as the center of the Gaussian, and the RF radius as the largest distance between points on the border of the 2D Gaussian at 1.5 standard deviations. To ensure high precision in all subsequent analyses, we excluded a small portion of neurons from all subsequent experiments whose Gaussian fit presented a normalized error above 0.2. All grating experiments in the macaque V1 model were conducted for balanced stimuli, spanning from -0.2 to 0.2 in the model input scale (obtained by z-scoring the training data). We collected responses to stimuli of 36 orientations spanning from 0 to 180 degrees, 36 different phases spanning from 0 to 360 degrees, and 25 spatial frequencies spanning from 1.1 to 8.0 cycles per degree of visual field. For each neuron, we selected the stimuli and responses corresponding to the phase of maximum response. For the size tuning experiment, we centered stimuli at the Sparse Noise RF positions and considered disks of radii spanning from 0 to 2.3 degrees. Considering the limited size of the input space of the model (2.67 degrees), stimuli corresponding to the largest radii values correspond to fullfield stimuli. We again tested multiple phase values and selected the responses corresponding to the maximally activating phase. The grating summation field was estimated as the first radius corresponding to 95% of maximal response.

The orientation contrast experiment was performed presenting to each neuron a center stimulus corresponding to GSF and a surround stimulus separated from the center disk by a moat of 0.23 degrees and reaching image borders. Iso-oriented surround stimuli matched all center grating parameters, ortho-oriented surround stimuli matched all center grating parameters except for orientation, shifted by 90 degrees.

#### MEI and surround optimization on macaque V1 neurons

Similarly to the analysis using the mouse V1 model, macaque V1 neuron MEIs were obtained by changing the pixels in input space to maximize neuronal response. The optimization procedure consisted in a Stochastic Gradient Descent (SGD) of 1000 steps, with step size of 10. To minimize artifacts, gradients where blurred with a sigma of 3 pixels. After each optimization step, MEI values were linearly scaled to have mean 0 and 0.05 standard deviation. The MEI mask was estimated by thresholding the MEI at 1.5 standard deviations above the mean (following the same algorithm used in the mouse analysis). Surround stimuli were obtained following the same algorithm used in the mouse analysis, optimizing the surround, corresponding to the region outside of the MEI mask, to obtain maximally exciting or maximally suppressing stimuli (surround mean=0, surround RMS contrast=0.1). The optimization algorithm parameters were consistent for the center and surround MEI optimizations.

#### Local patches Gabor-fit analysis

We performed a quantitative analysis based on fitting Gabor functions to local patches extracted from the optimized surround images to quantify pattern completion of Gabor patterns in the excitatory and inhibitory surround images. In this analysis, we only used neurons whose MEI was well fitted by a Gabor function (normalized fit error threshold = 0.2, spatial frequency threshold = 1, number of remaining neurons=126). We extracted local patches *I*_*patch*_ = *M* ∗*I* from optimized surround images *I* using a truncated isotropic Gaussian mask 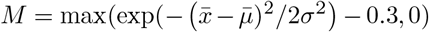 .*σ* was set to be 0.34 degrees of visual angle and the mask was placed in 4 cardinal positions (with respect to preferred orientation) for each neuron considered. Specifically, the centers of the local patches were placed at neuron dependent distance corresponding to the size of the MEI mask in the direction considered (2 collinear direction, 2 orthogonal direction). In this way, we ensured that the local patch was encompassing a significant part of MEI and surround. During the fit, we restrain some parameters to ensure that the resulting Gabor corresponded to an oriented feature extracting pattern (aspect ratio < 1.5 and spatial frequency > 1 cycle per degree). To distinguish between good and poor fits, we selected a normalized fit error threshold of 0.3.

#### Divisive normalization model

We considered a population of 10,000 LN Gabor filter simple cells of randomly sampled orientation, position and phase. Gabor filter parameters considered are: spatial frequency of 2.5, *σ* of 0.2, aspect ratio of 1, image resolution of 93x93 (2.67×2.67 degrees). These parameters generate Gabors filters resembling the MEI of the macaque V1 neurons. We use ReLU to enforce non-negative responses. We then implemented a divisive normalization model (see Heeger (1992)):

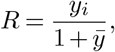

where 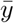 represents the response of the population, divisively normalizing the response ^*yi*^ of another simple cell *i*. We used ELU() + 1 as nonlinearity of neuron *i* to allow gradient flow during optimization. We obtained the MEI of neuron of the Heeger model neuron by optimization the input space to elicit maximal response. We enforced pixel mean to 0, pixel standard deviation to 0.05, and trained using SGD with step size of 0.1, 1000 steps and gradient Gaussian blurring of 1 pixel. We enforced pixel mean to 0, pixel standard deviation to 0.05, and trained using SGD with step size of 0.1, 1000 steps and gradient Gaussian blurring of 1 pixel. We identified a MEI mask (threshold=1.5) and optimized the surround to maximally suppress MEI response. In this case we enforced pixel mean to 0, pixel standard deviation to 0.10, and trained using SGD with step size of 0.1, 3000 steps and gradient Gaussian blurring of 1 pixel.

#### Replication of center-surround modulation in functional connectomics dataset

Recently, we and others released a large-scale functional connectomics dataset of mouse visual cortex (“MICrONS dataset”), including responses of >75k neurons to full-field natural movies and the reconstructed sub-cellular connectivity of the same cells from electron microscopy data (MICrONS Consortium et al., 2021). A dynamic recurrent neural network (RNN) model of this mouse’s visual cortex—digital twin—exhibits not only a high predictive performance for natural movies, but also accurate out-of-domain performance on other stimulus classes such as drifting Gabor filters, directional pink noise, and random dot kinematograms (Wang et al., 2023). Here, we took advantage of the model’s ability to generalize to other visual stimulus domains and presented our full-field and masked images to this digital twin model in order to relate specific functional properties to the neurons’ connectivity and anatomical properties. Specifically, we recorded the visual activity of the same neuronal population to static natural images as well as to the identical natural movies that were used in the MICrONS dataset. Neurons were matched anatomically as described for the closed loop experiments. Based on the responses to static natural images we trained a static model as described above, and from the responses to natural movies we trained a dynamic model using a RNN architecture described in (Wang et al., 2023). This enabled us to compare the MEIs and surround images for the same neurons generated from two different static models: one trained directly on responses from real neurons, and another trained on synthetic responses to static images from dynamic models. We then presented MEIs and optimized surround images to the animal in a closed-loop experiment.

To investigate the circuitry implementation of pattern completion, we combined synaptic connectivity data extracted from electron microscopy imaging with functional tuning data obtained from the digital twin model. Receptive field overlap between pairs of neurons was quantified using the intersection over union (IoU) of their MEI masks. Additionally, feature tuning similarity between neurons was assessed using the digital twin model, which comprises a shared core for visual feature extraction and a final readout layer where the extracted visual features are linearly weighted to predict neuronal activity. The feature similarity between pairs of neurons is measured as the cosine similarity of their feature weights. Neurons with reliable visual responses (*CC*_*max*_ *>* 0.4) that are well predicted by the digital twin model (*CC*_*abs*_ *>* 0.2) were included in the downstream analysis. Visual response reliability (*CC*_*max*_) and model performance (*CC*_*abs*_) were quantified as described in (Wang et al., 2023).

We conducted Welch’s t-test to compare feature similarity between connected neurons and randomly paired unconnected neurons at different levels of receptive field overlap. Corrections for multiple comparisons were applied using the Benjamini–Hochberg procedure.

To further examine the relationship between feature similarity and connectivity, we modeled the number of synapses between neuron pairs (*n*_*syn*_) using a Poisson generalized linear model of form:

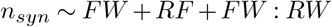

This model incorporated feature similarity (FW), receptive field overlap (RF), and their interaction term (FW:RF). The Likelihood Ratio Test (LRT) was employed to assess whether the inclusion of the interaction term significantly improved model fit compared to a reduced model without it.

### Probabilistic model

#### Generative model

Our generative model is hierarchical and probabilistic, containing three groups of random variables: **g, x** and **I. g***∈* {0, 1}^*N*^ represents the presence *N* high level textures and objects, modeled as independent Bernoulli distributions:

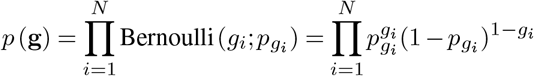

where *p*_*gi*_ is the *a priori* probability that the feature represented by *g*_*i*_ is present in the image.

**x***∈*{0, 1}^9*×N*^ represents the presence of 9*×N* local visual features, modeled as a Bernoulli distribution conditioned on **g**:

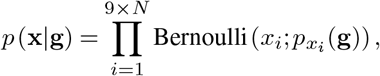

where *p*_*xi*_ (**g**) represents the prior expectation of whether feature *x*_*i*_ is present given the presence of the global features represented by **g**. Specifically, those elements of **x** representing local features compatible with the presence of any one *g*_*i*_ have a high probability, *p*_high_, when *g*_*i*_ = 1, and otherwise a low probability, *p*_low_. We assign *p*_high_ to be 0.80 and *p*_low_ to be 0.02 but our qualitative results do not depend on the specific values. **I***∈* ℝ^*H×W*^ represents the image of height *H* and width *W*, and is a modeled as the linear combination of the projective fields (PFs) inferred to be present in the image, corrupted by isotropic Gaussian pixel noise of variance *σ*^2^ (Olshausen and Field, 1996):

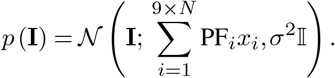

Note that the RF of a neuron is closely related to the PF but slightly different (Olshausen and Field, 1996).

#### Inference

We assume the neural responses are proportional to the marginal posterior probabilities, *p*(*x*_*i*_|**I**), of the elements of **x** each representing a different V1 neuron (but note that any monotonic relationship will yield the same qualitative results). We compute the posterior for various input images using Python’s PyMC package (Oriol et al., 2023) to obtain the average simulated responses (stars in Figure 5f,g). In order to simulate trial-to-trial variability, we interpret (binary) samples as spikes (Buesing et al., 2011) and compute the per trial firing rates in Figure 5f,g by counting the number of spikes over a trial duration of 1s assuming a sampling rate of 1/20ms.

## Code and data availability

Our coding framework uses general tools like PyTorch, Numpy, scikit-image, matplotlib, seaborn, DataJoint (Yatsenko et al., 2015, 2018, 2021), Jupyter, and Docker. All custom analysis code and all data will be publicly available in an online repository latest upon journal publication. Please contact us if you would like access before that time.

## ACKNOWLEDGEMENTS

The authors thank David Markowitz, the IARPA MICrONS Program Manager, who coordinated this work during all three phases of the MICrONS program. We thank IARPA program managers Jacob Vogelstein and David Markowitz for co-developing the MICrONS program. We thank Jennifer Wang, IARPA SETA for her assistance.

The work was supported by the Intelligence Advanced Research Projects Activity (IARPA) via Department of Interior/ Interior Business Center (DoI/IBC) contract numbers D16PC00003, D16PC00004, and D16PC0005. The U.S. Government is authorized to reproduce and distribute reprints for Governmental purposes notwithstanding any copyright annotation thereon. XP acknowledges support from NSF CAREER grant IOS-1552868. XP and AST acknowledge support from NSF NeuroNex grant 1707400. AST also acknowledges support from the National Institute of Mental Health and National Institute of Neurological Disorders And Stroke under Award Number U19MH114830 and National Eye Institute award numbers R01 EY026927 and Core Grant for Vision Research T32-EY-002520-37. RMH acknowledges support from NSF/CAREER IIS-2143440 and National Eye Institute R01 EY028811. Disclaimer: The views and conclusions contained herein are those of the authors and should not be interpreted as necessarily representing the official policies or endorsements, either expressed or implied, of IARPA, DoI/IBC, or the U.S. Government. This work was also supported by the German Research Foundation (to FHS, KF & SS: SFB 1233, Robust Vision: Inference Principles and Neural Mechanisms, TP06 and TP11, project number 276693517; to ASE & FHS: SFB 1456, project B05 & A06, project number 432680300), the European Research Council (ERC) under the European Union’s Horizon Europe research and innovation programme (grant agreement number 101041669), and the ERDF-Project Brain dynamics 1180 (CZ.02.01.01-00-22-008-0004643). PAP was funded through a ZIM grant by the German Federal Ministry for Economic Affairs and Energy (ZF4076506AW9).

## AUTHOR CONTRIBUTIONS

**JK**: Conceptualization, Methodology, Validation, Software, Formal Analysis, Investigation, Writing - Original Draft, Visualization, Project administration; **SS**: Conceptualization, Formal Analysis, Software, Writing - Original Draft, Visualization; **LB**: Conceptualization, Formal Analysis, Software, Writing - Original Draft, Visualization; **KP, TM, RF, LN**: Investigation, Validation; **ZhuD, EW**: Investigation, Validation, Methodology (dynamic model and functional connectomics); **ZhiD, DTT**: Conceptualization, Methodology, Visualization (dynamic model and functional connectomics); **PGF, StP, SaP, JR**: Investigation, Validation, Methodology (functional connectomics); **KFW**: Methodology, Formal Analysis, Software, Writing - Review & Editing; **ASE, XP**: Conceptualization, Writing - Review & Editing, Funding acquisition; **PAP**: Software, Methodology, Formal Analysis, Writing - Review & Editing; **JA**: Conceptualization, Writing - Review & Editing, Funding acquisition, Supervision; **RMH**: Conceptualization, Methodology, Supervision, Funding acquisition, Writing - Review & Editing; **FHS**: Conceptualization, Methodology, Writing - Review & Editing, Supervision, Funding acquisition; **AST**: Conceptualization, Experimental and analysis design, Supervision, Funding acquisition, Writing - Review & Editing, Project administration; **KF**: Conceptualization, Formal Analysis, Supervision, Visualization, Writing - Original draft, Project administration, Supervision, Funding acquisition. Fu *et al*. | Pattern completion and disruption characterize contextual modulation in the visual cortex

## Supplementary Information

**Supplemental Fig. 1.**
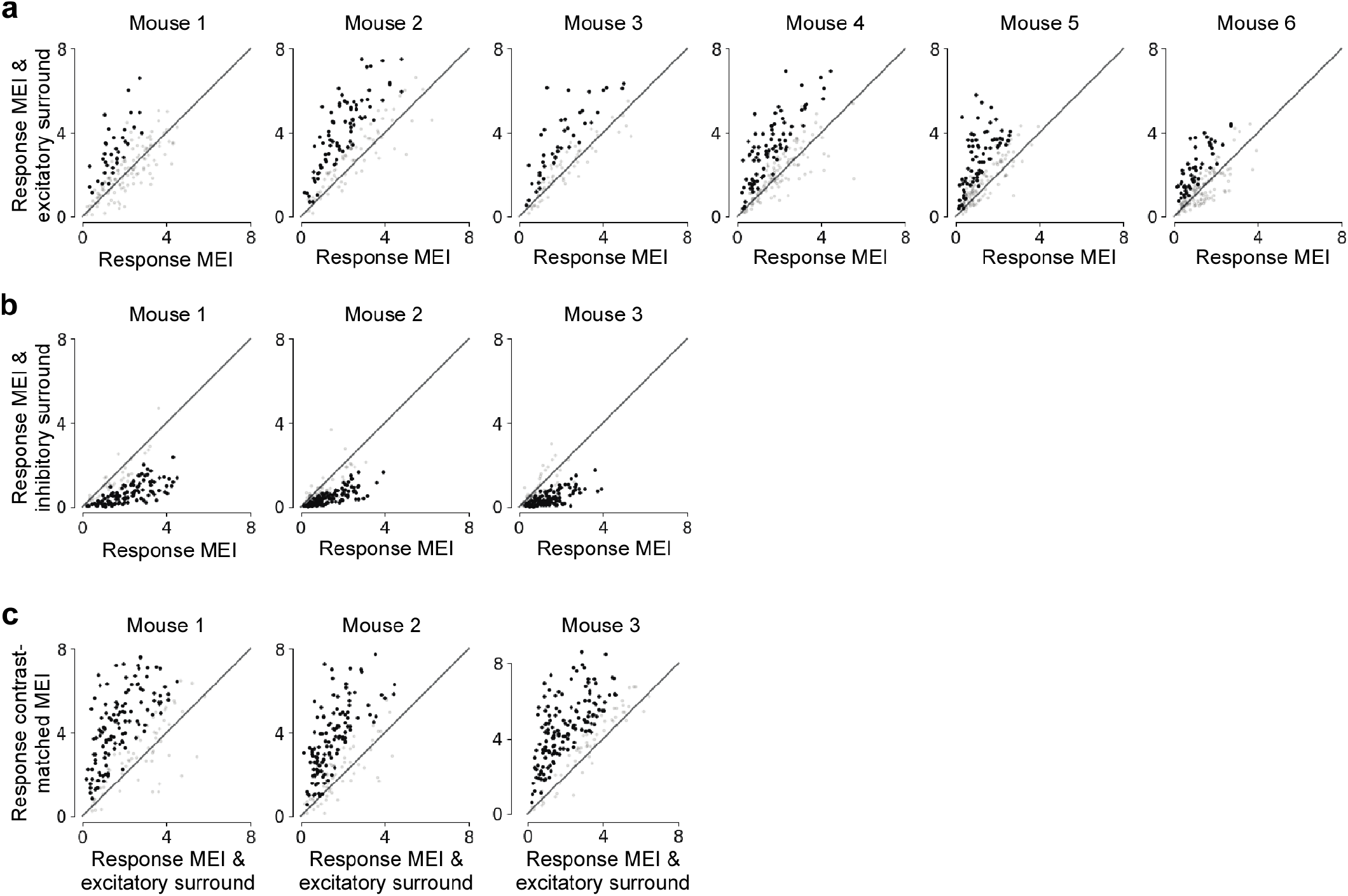
Neuronal responses to MEIs and surround imaged recorded during inception loop experiments across animals. **a**, Comparing observed responses to the MEI (x-axis) and the excitatory surround (y-axis) per experiment (n=6 mice, 960 cells total). Dark dots indicate neurons where the response to the surround images is significantly higher than to the MEI (Wilcoxon rank-sum test, p-values<0.05). Across the population, the modulation was significant for all animals (p-values<0.05, Wilcoxon signed rank test). **b**, Comparing observed responses to the MEI (x-axis) and the inhibitory surround (y-axis) per experiment (n=3 mice, 510 cells total). Dark dots indicate neurons where the response to the surround images is significantly lower than to the MEI (Wilcoxon rank-sum test, p-value<0.05). Across the population, the modulation was significant for all animals (p-value<0.05, Wilcoxon signed rank test). **c**, Comparing observed responses to the excitatory surround (x-axis) and the contrast-matched MEI (y-axis) per experiment (n=3 mice, 560 cells total). Dark dots indicate neurons where the response to the contrast-matched MEIs is significantly higher than to the MEI (Wilcoxon rank-sum test, p-value<0.05). Across the population, the modulation was significant for all animals (p-value<0.05, Wilcoxon signed rank test).

**Supplemental Fig. 2.**
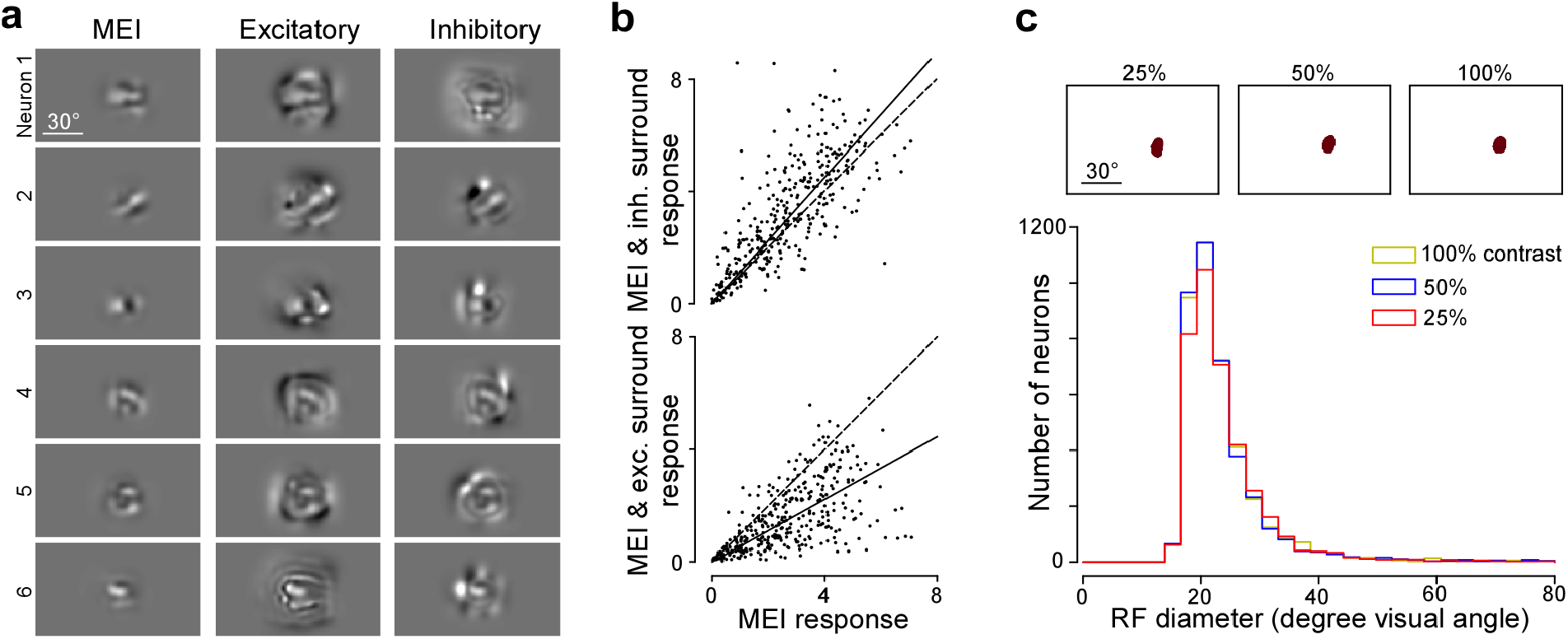
Center-surround effects are preserved at higher contrast. **a**, MEI and MEI with excitatory and inhibitory surround for six example neurons, optimized with a higher contrast constraint (0.1 for the MEI (instead of 0.05) and 0.2 for the surround (instead of 0.1)). **b**, Recorded neuronal responses to MEI and MEI with excitatory and inhibitory surround. Neuronal responses to the MEI were significantly modulated (p-value=4. × 10^*−*13^, 0.00949, 3.20 × 10^*−*05^, for excitatory surround, p-value=1.53 × 10^*−*22^, 5.85 × 10^*−*18^, 7.12 × 10^*−*18^ for inhibitory surround, Wilcoxon signed-rank test). **c**, Distribution of RF diameters estimated using a sparse noise stimulus with different contrast levels. The change is RF sizes across different contrast levels is minimal (100%*vs*50% : *−*0.009 ± 8.87, 50%*vs*25% : *−*0.52 ± 9.66, 100%*vs*25% : *−*0.53 ± 9.31, mean±std in degrees of visual angle).

**Supplemental Fig. 3.**
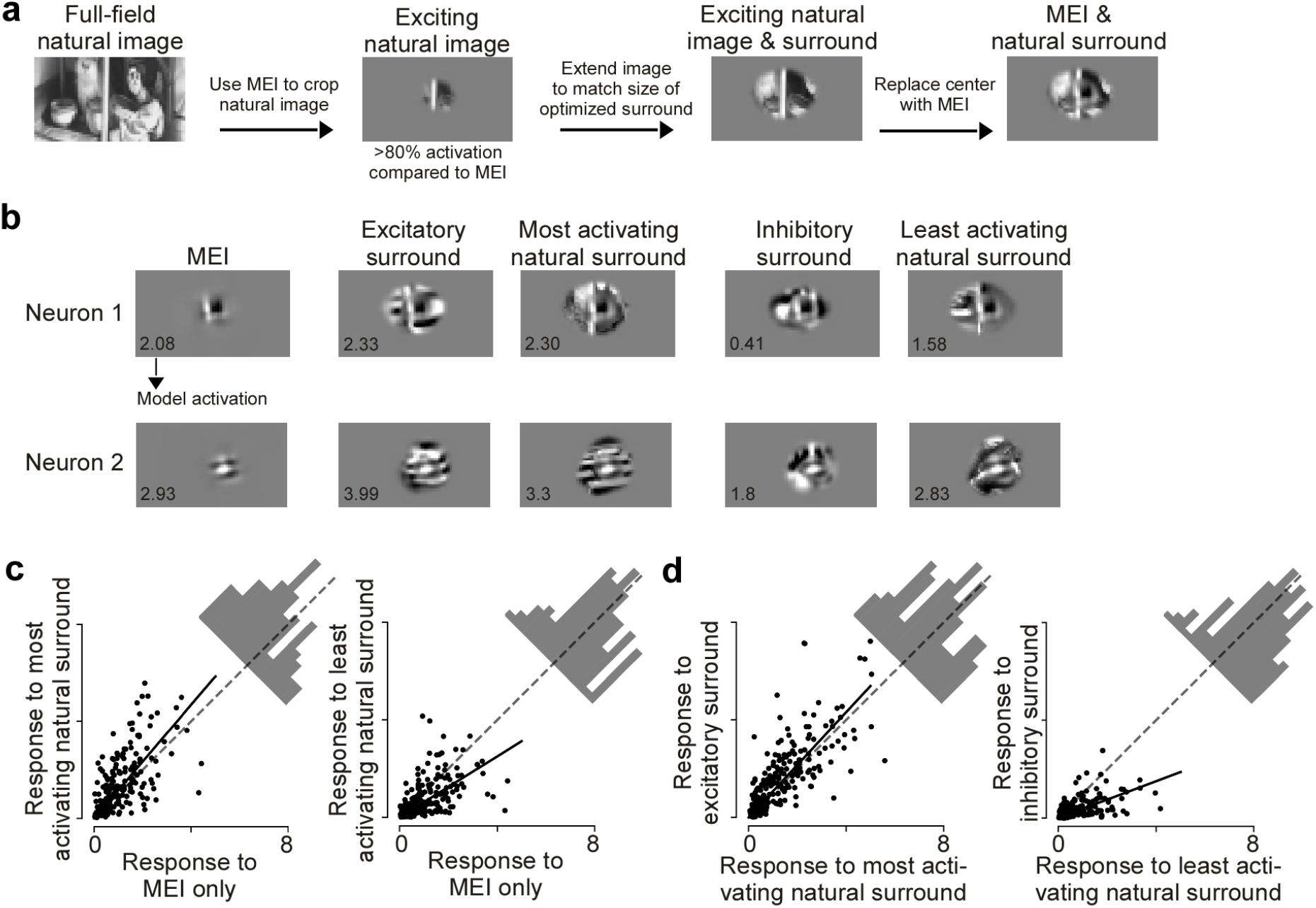
Surround images correspond to the optimal modulating stimulus and are ecologically relevant. **a**, Schematic illustrating how we obtained natural surround images for one example neuron. **b**, Optimized excitatory and inhibitory surround images, most exciting and inhibiting natural surrounds and MEI of two example neurons. The predicted activation score is indicated in the bottom left of the images. **c**, Observed responses to the MEI with natural surround images compared to the MEI alone. Across the population, the least activating natural surround images suppressed neuronal response (p-value=1.84 × 10^*−*8^, Wilcoxon signed rank test), and the most activating natural surround images enhanced neuronal response (p-value=2.44 × 10^*−*9^, Wilcoxon signed rank test). Across stimulus repetitions, 23% responded significantly stronger to the most activating natural images than to the MEI (n=3 animals, 226 cells, two-sided t-test, p-value<0.05) and 25% of the neurons responded significantly weaker to the least activating natural surround images than to the MEI. Solid line indicates the regression line across the population, and dotted gray line indicates the diagonal. **d**, Observed responses to the MEI with natural surround images compared to the MEI with excitatory/inhibitory surround. Across the population, the MEI with inhibitory surround suppressed neuronal response more than the MEI with the least activating natural surround (p-value=1.98 × 10^*−*20^, Wilcoxon signed rank test). The MEI with excitatory surround enhanced neuronal response more than the MEI with most activating natural surround (p-value=1.05 × 10^*−*6^, Wilcoxon signed rank test). Across stimulus repetitions, 37% of neurons responded significantly weaker to the MEI with inhibitory surround compared to the MEI with the least activating natural surround and 19% of the neurons responded significantly stronger to the MEI with excitatory surround compared to the MEI with the most activating natural surround (n=3 animals, 226 cells, two-sided t-test, p-value<0.05). Solid line indicates the regression line across the population, and dotted gray line indicates the diagonal. Please note that due to spatial correlations in natural images, spatial patterns of inhibitory surround MEIs characterized by disruption are less common in natural scenes. Should we screen a larger pool of natural images (e.g., 1 million), we anticipate observing natural surrounds that inhibit as effectively as optimized surrounds.

**Supplemental Fig. 4.**
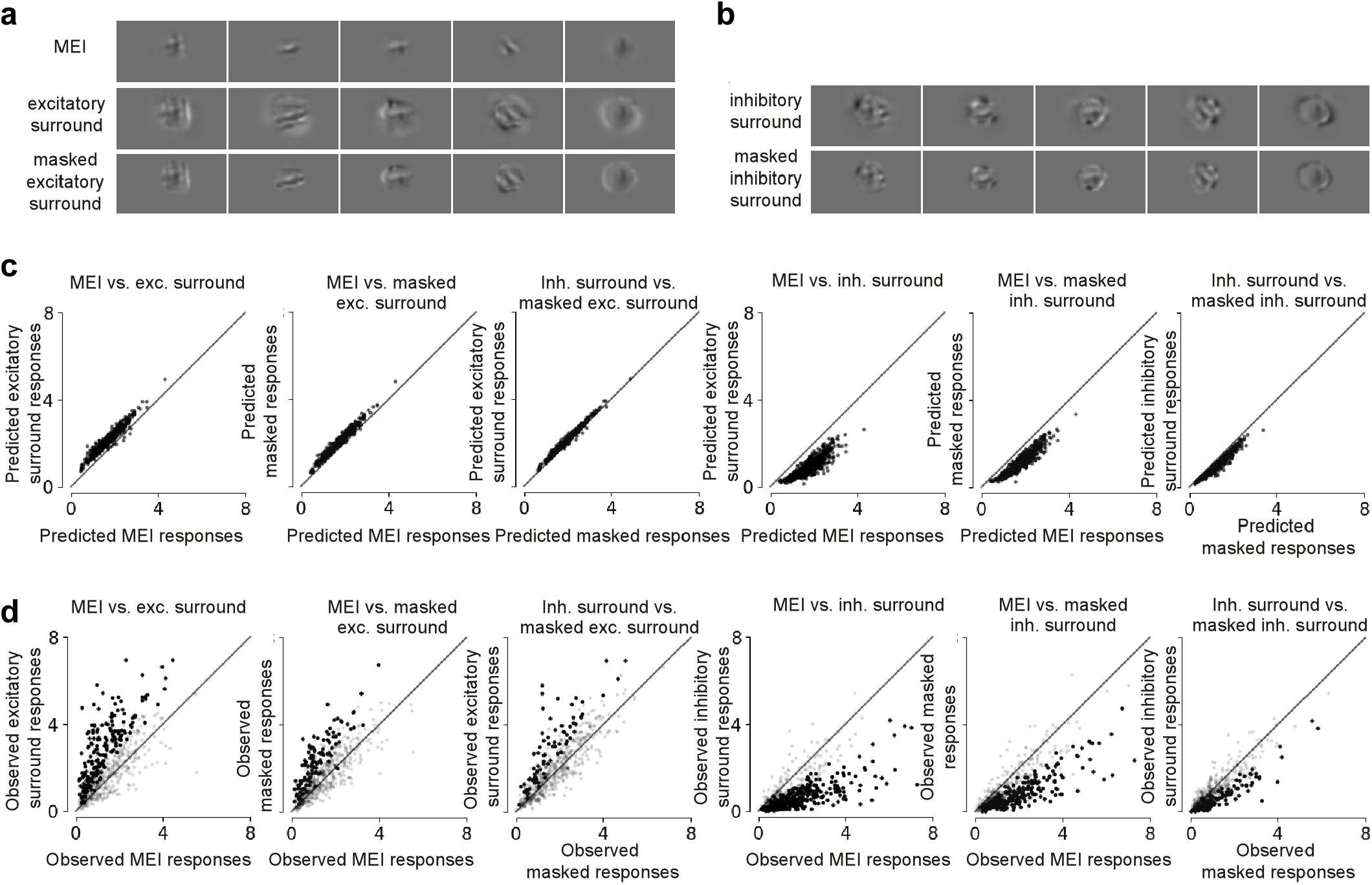
Images restricted to the far surround still result in surround modulation. **a**, Examples of the MEI, the excitatory surround and cropped excitatory surround. **b**, Examples of the MEI, the inhibitory surround and cropped inhibitory surround. **c**, Comparing predicted response to the MEI, the excitatory surround and the cropped surround image (n=3, 560 cells). **d**, Comparing predicted response to the MEI, the inhibitory surround and the cropped surround image (n=3, 560 cells). **e**, Comparing observed response to the MEI, the excitatory surround and the cropped surround image (n=3, 560 cells). Black dots indicate neurons with significantly higher response under the condition on the y-axis (one-sided Wilcoxon rank-sum test, p<0.05, 33.6%, 20.2% and 13.4% significant cells for each pair). Modulation is significant on population level for each pair (p-value=1.83 × 10^*−*45^, 9.98 × 10^*−*45^, 6.89 × 10^*−*19^, Wilcoxon signed rank test). **f**, Comparing observed response to the MEI, the inhibitory surround and the cropped surround image (n=3, 560 cells). Black dots indicate neurons with significantly higher response under the condition on the y-axis (one-sided Wilcoxon rank-sum test, p<0.05, 55.9%, 40.3% and 19.6% significant cells for each pair). Modulation is significant on population level for each pair (p-value=8.05 × 10^*−*73^, 9.03 × 10^*−*66^, 2.42 × 10^*−*24^, Wilcoxon signed rank test).

**Supplemental Fig. 5.**
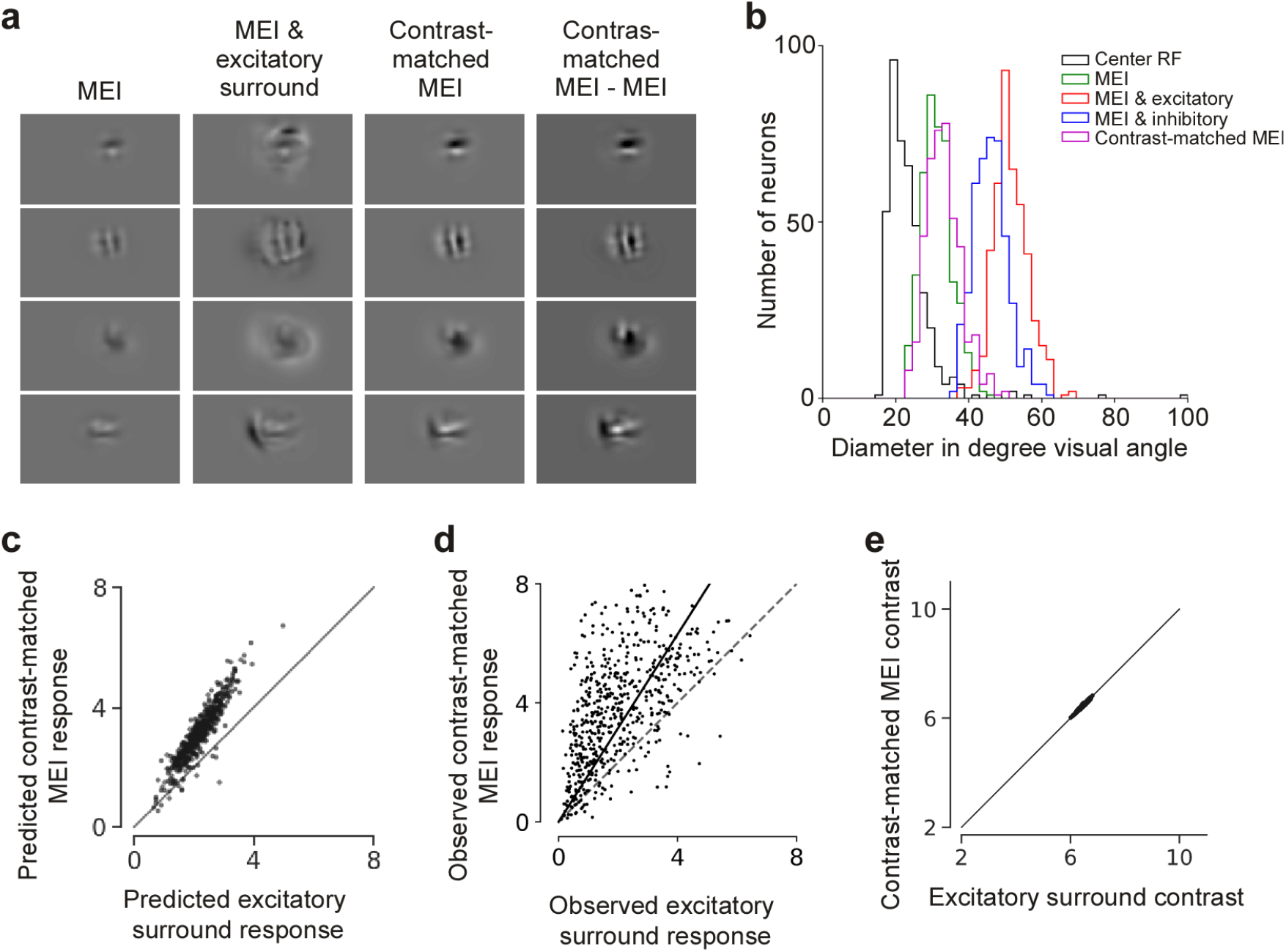
Contrast-matched MEIs result in higher activation than MEIs with excitatory surround. **a**, Panel shows MEI, excitatory surround with MEI, the contrast-matched MEI, and the difference between the original MEI and the contrast-matched MEI for 4 example neurons. Note that the contrast-matched MEI is a scaled-up version of the original MEI with same features. **b**, Diameters of RFs estimated using sparse noise, the MEIs, the MEIs with excitatory and inhibitory surround, and the contrast-matched MEI. Same data shown in Fig. 2e except for the contrast-matched MEI. The mean of the contrast-matched MEI (magenta distribution) size across all neurons (n=4, 434 cells) is 33.2 degrees ± 0.23 (mean ± s.e.m.). The size of the contrast-matched MEI is slightly larger than the original MEI (31.3 degrees ± 0.20). **c**, Model predicted responses to the MEI and excitatory surround (x-axis) and contrast-matched MEI (y-axis). Responses are depicted in arbitrary units, corresponding to the output of the model. **d**, Observed responses to the the MEI and excitatory surround (x-axis) and contrast-matched MEI (y-axis). For each neuron, responses are normalized by the standard deviation of responses to all images. Across the population, the neuronal responses to the contrast-matched MEI was significantly higher (p-value=7.35 × 10^*−*80^, Wilcoxon signed rank test, slope of linear regression line=1.58). Across stimulus repetitions, 58.9% of the neurons responded stronger to the contrast-matched MEI (n=3 animals, 560 cells, two-sided t-test, p-value<0.05). Solid line indicates the regression line across the population, and dotted gray line indicates the diagonal. **e**, Contrast comparison between the MEI and excitatory surround (x-axis) and the contrast-matched MEI. By definition, the full-field contrast of each pair of images are matched.

**Supplemental Fig. 6.**
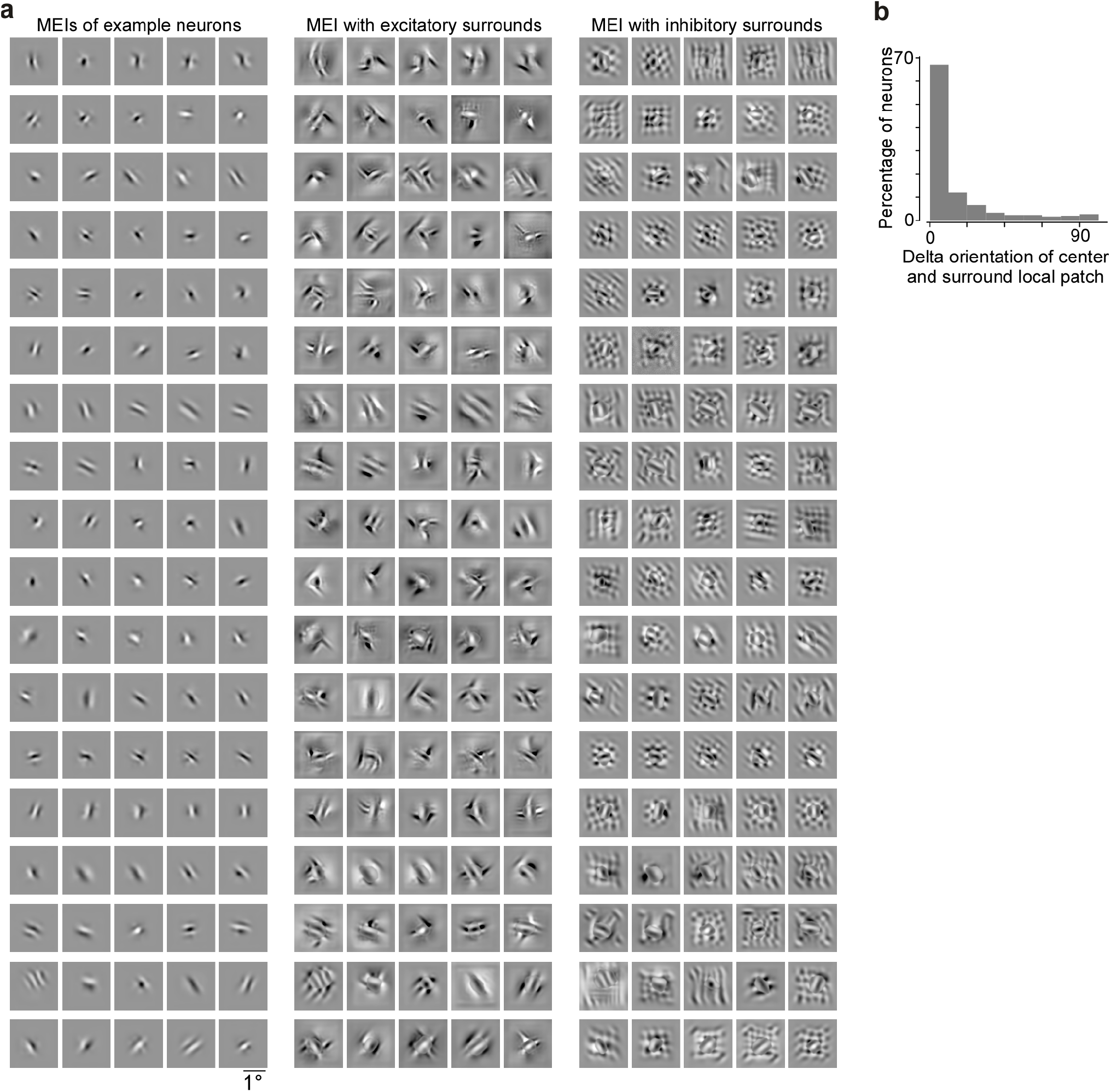
MEIs with excitatory and inhibitory surrounds of macaque V1 neurons. **a**, MEIs of example neurons (left) used for the collinearity analysis shown in Fig. 5, with excitatory and inhibitory surround images optimized through the model. Order of neurons matches across the three columns. **b**, Distribution of difference in orientation (delta orientation) between center preferred orientation and orientation of Gabor fit to surround local patch.

